# Experience shapes activity dynamics and stimulus coding of VIP inhibitory and excitatory cells in visual cortex

**DOI:** 10.1101/686063

**Authors:** Marina E. Garrett, Sahar Manavi, Kate Roll, Douglas R. Ollerenshaw, Peter A. Groblewski, Justin Kiggins, Xiaoxuan Jia, Linzy Casal, Kyla Mace, Ali Williford, Arielle Leon, Stefan Mihalas, Shawn R. Olsen

## Abstract

Cortical circuits are flexible and can change with experience and learning. However, the effects of experience on specific cell types, including distinct inhibitory types, are not well understood. Here we investigated how excitatory and VIP inhibitory cells in layer 2/3 of mouse visual cortex were impacted by visual experience in the context of a behavioral task. Mice learned to perform an image change detection task with a set of eight natural scene images, viewing these images thousands of times. Subsequently, during 2-photon imaging experiments, mice performed the task with these familiar images and three additional sets of novel images. Novel images evoked stronger overall activity in both excitatory and VIP populations, and familiar images were more sparsely coded by excitatory cells. The temporal dynamics of VIP activity differed markedly between novel and familiar images: VIP cells were stimulus-driven by novel images but displayed ramping activity during the inter-stimulus interval for familiar images. Moreover, when a familiar stimulus was omitted from an expected sequence, VIP cells showed extended ramping activity until the subsequent image presentation. This prominent shift in response dynamics suggests that VIP cells may adopt different modes of processing during familiar versus novel conditions.

**HIGHLIGHTS:**

- Experience with natural images in a change detection task reduces overall activity of cortical excitatory and VIP inhibitory cells
- Encoding of natural images is sharpened with experience in excitatory neurons
- VIP cells are stimulus-driven by novel images but show pre-stimulus ramping for familiar images
- VIP cells show strong ramping activity during the omission of an expected stimulus

## INTRODUCTION

Neural circuits are dynamically shaped by experience and expectation (De Lange et al., 2018; LeMessurier and Feldman, 2018; Ranganath and Rainer, 2003). Visual experience can produce modifications of cortical representations, including changes in response gain, selectivity, correlations, and population dynamics (Jurjut et al., 2017; Khan et al., 2018; Makino and Komiyama, 2015; Poort et al., 2015; Weskelblatt and Niell, 2019; Woloszyn and Sheinberg, 2012). Moreover, sensory and behavioral experience can lead to the emergence of predictive activity in the visual cortex including reward anticipation (Poort et al., 2015; Shuler and Bear, 2006), spatial expectation (Fiser et al., 2016; Saleem et al., 2018), pattern completion (Gavornik and Bear, 2014), and prediction error signals (Fiser et al., 2016; Hamm and Yuste, 2016; Homann et al., 2017). Visual cortical circuits are also influenced by behavioral context, internal states, and other factors beyond external stimulus features (Batista-Brito et al., 2018; Busse et al., 2017; Gilbert and Li, 2013; Khan and Hofer, 2018; Kuchibhotla and Bathellier, 2018; McGinley et al., 2015; Pakan et al., 2018). These experience and state-dependent changes in sensory cortex can involve top-down feedback (Fiser et al., 2016; Makino and Komiyama, 2015; Petro et al., 2014; Zhang and Dan, 2014) and neuromodulatory inputs (Chubykin et al., 2013; Kuchibhotla et al., 2017; Pinto et al., 2013), and may be associated with a shift in the balance of bottom-up sensory and top-down contextual signals conveying internal states and learned expectations. Inhibitory interneurons are likely to play a key role in this process by dynamically regulating the flow of information (Chiu et al., 2019; Hangya et al., 2014; Kepecs and Fishell, 2014). Elucidating how different cell populations, particularly inhibitory cells, contribute to experience-dependent changes in sensory coding is critical to understand the dynamic nature of cortical circuits.

Vasoactive intestinal peptide (VIP) expressing cells comprise a major class of inhibitory neurons and are well-positioned to mediate top-down influences on local circuits in sensory cortex. VIP cells receive long-range projections from frontal areas (Lee et al., 2013; Wall et al., 2016; Zhang and Dan, 2014) as well as cholinergic and noradrenergic inputs (Alitto and Dan, 2013; Fu et al., 2014). VIP cells are highly active during states of arousal (Fu et al., 2014; Reimer et al., 2014), are modulated by task engagement (Kuchibhotla et al., 2017), and are responsive to behavioral reinforcement (Pi et al., 2013). In the local cortical circuitry, VIP cells primarily inhibit another major class of inhibitory interneuron, somatostatin (SST) cells (Lee et al., 2013; Munoz et al., 2017; Pfeffer et al., 2013; Pi et al., 2013), which can result in disinhibition of excitatory neurons (Fu et al., 2017; Lee et al., 2013; Letzkus et al., 2011). SST cells target the apical dendrites of pyramidal neurons (Kepecs and Fishell, 2014) and removal of this inhibition may facilitate the association of top-down and bottom-up input by pyramidal cells (Chen et al., 2015; Larkum, 2012; Makino and Komiyama, 2015). However, little is known about how VIP cell activity is modified by visual experience. As VIP cells exhibit strong surround suppression in response to high contrast oriented gratings (Dipoppa et al., 2018), the use of naturalistic stimuli may be critical to uncover the role of VIP cells in regulating the flow of information with experience.

Here we investigated how long-term behavioral experience with natural scene images alters activity of cortical excitatory and VIP inhibitory cells in layers 2/3 of mouse visual cortex. Mice were trained to perform a change detection task in which images are flashed in a periodic manner and mice were rewarded for detecting changes in image identity. Mice learned the task with one set of eight natural images, which were viewed thousands of times during training and were thus highly familiar. During subsequent 2-photon imaging, these familiar images as well as three novel image sets were tested. Familiar images were associated with lower overall population activity in both excitatory and VIP cells, and excitatory cells were more stimulus selective for familiar images. Notably, VIP inhibitory cells had distinct activity dynamics during sessions with familiar versus novel images. VIP cells were stimulus-driven by novel images but displayed ramping activity between stimulus flashes when presented with familiar images. These cells showed even greater activity when an expected stimulus was omitted from the regular image sequence. Overall, these results demonstrate experience-dependent changes in across two cortical cell classes and suggest that VIP cells may adopt different modes of processing during familiar versus novel conditions.

## RESULTS

### Visual change detection task with familiar and novel images

We trained mice to perform a go/no-go visual change detection task with natural scene stimuli. In this task, mice are presented with a continuous stream of flashing images (Figure 1A-C). On ‘go’ trials, a change in image identity occurs at a time unknown to the animal (Figure 1B). To earn water rewards, mice must report the image change by licking a reward spout within a 750 ms response window. False alarms are quantified during ‘catch’ trials in which the repeated image does not change, but licking behavior is measured in a similar time window. To test whether expectation signals were present in the visual cortex during this task, we randomly omitted ∼5% of all non-change flashes; this appears as an extended gray period to the mouse and corresponds to a gap in the regular timing of stimuli (Figure 1C).

**Figure 1:**
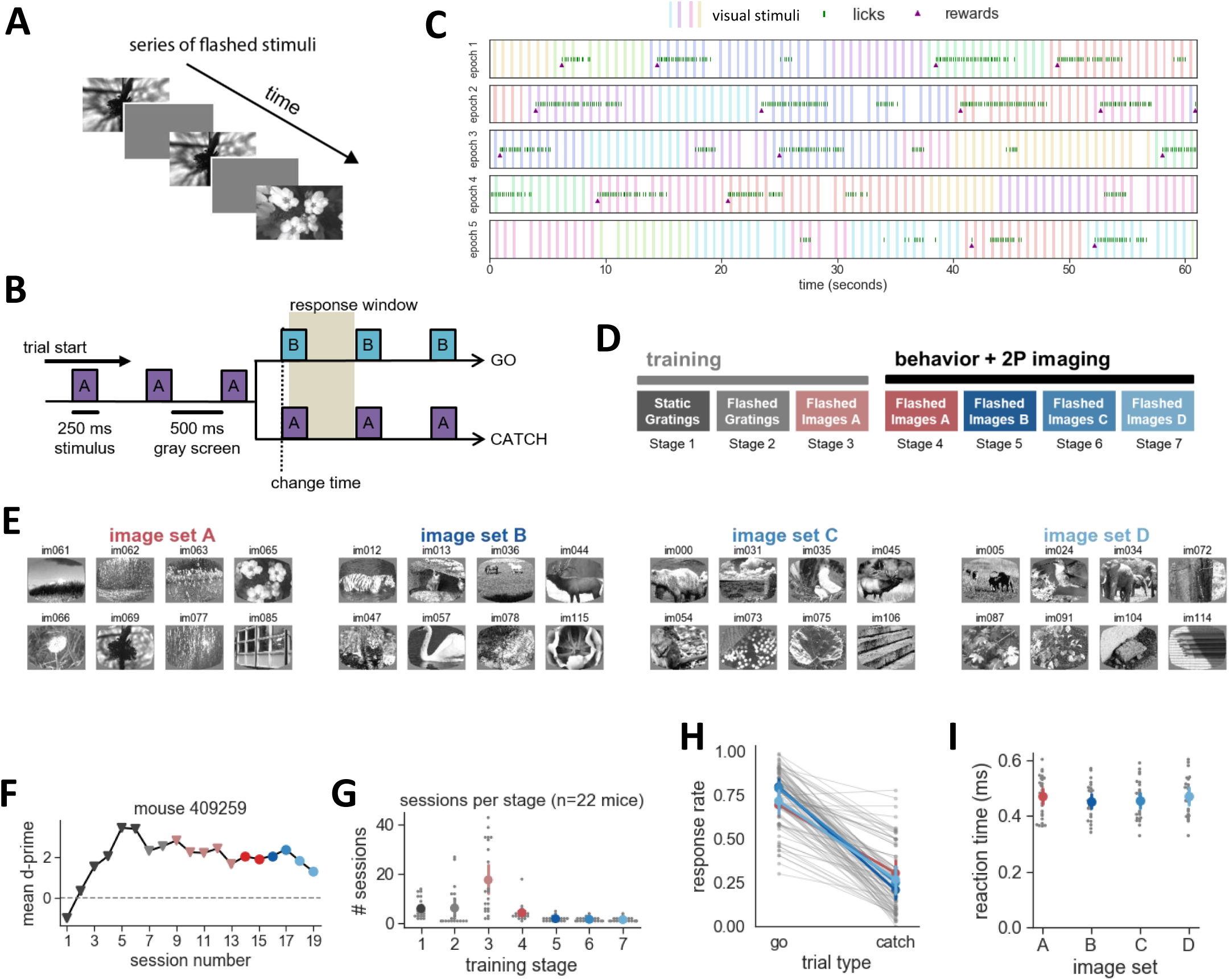
Natural scene change detection task with familiar and novel images. **A**) Schematic of stimulus presentation during the task. Images are presented for 250ms followed by 500ms of gray screen. **B)** Schematic of trial structure. On go trials, a change in image identity occurs and mice must lick within the 750ms response window to receive a water reward. On catch trials, no stimulus change occurs and the behavioral response is measured to quantify guessing behavior. **C)** Example behavior performance across 5 minutes of one session, separated into 5 one-minute epochs. Colored vertical bars indicate stimulus presentations, different colors are different images. Licks are shown as green tick marks. Licks within 750ms of a change result in reward delivery, shown as purple triangles. 5% of all non-change image flashes are omitted, visible as a gap in the otherwise periodic stimulus sequence. **D)** Training sequence. Mice are initially trained with static gratings of 2 orientations, first with no gray screen in between grating presentations (stage 1), then introducing the 500ms inter stimulus interval (stage 2). Next, mice are transitioned to change detection with 8 natural scene images (stage 3, image set A). During the 2-photon imaging portion of the experiment, mice are tested with image set A as well as 3 novel image sets (B, C, D) on subsequent days. **E)** The four sets of 8 natural images were shown, in separate sessions. **F)** Example training trajectory of one mouse. Triangles represent behavior training and circles represent behavior + 2P imaging sessions. **G)** Number of sessions spent in each stage across mice. Mean +/- 95% confidence intervals in color, individual mice in gray. **H)** Response rates for go and catch trials are similar across image sets, demonstrating that mice generalize task performance to novel images. Individual behavior sessions are shown in gray and average +/- 95% confidence intervals across sessions for each image set are shown in colors corresponding to image sets in C (p>0.05 for all image set comparisons). **I)** No significant difference in response latency across image sets (p>0.05 for all image set comparisons). Mean +/- 95% confidence intervals in color, individual sessions in gray.

Behavioral training proceeded through a series of stages: mice first learned task rules with full-field gratings of two different orientations and no intervening gray period, followed by introduction of a 500ms gray screen period between flashes, then training with natural scene images, and finally simultaneous behavior and 2-photon imaging with multiple image sets, where omitted flashes were introduced (Figure 1D). During the natural image training stage (stage 3), mice were trained with one set of eight images (image set A) for an extended number of sessions (range = 6-46 sessions with image set A, median = 17 sessions; Figure 1G, Supplemental Figure 1A). On average, mice viewed each of the eight scenes from the familiar image set 10,350 times prior to the 2-photon imaging stage (range: 944-26,784 individual stimulus presentations per image).

During the physiology portion of the experiment, mice were challenged with either the familiar image set or one of three additional novel image sets (Figure 1E,F). Mice performed the task with similar hit and false alarm rates across image sets (Figure 1H, Supplemental Figure 1B,C; statistics are described in the figure legends throughout). Reaction times were also similar for familiar and novel image sets (Figure 1I). During the task, mice are free to run on a circular disk and typically stop running when emitting a licking response. There was no difference in average running speed or pattern of running behavior during novel and familiar image sessions (Supplemental Figure 1D,E). Together, these results indicate that overall behavior was similar for novel and familiar image sets.

### Imaging excitatory and inhibitory cortical populations during task performance

We measured activity in transgenic mice expressing the calcium indicator GCaMP6f in excitatory pyramidal neurons (Slc17a7-IRES2-Cre; CaMKII-tTA; Ai93-GCaMP6f) or VIP inhibitory neurons (VIP-IRES-Cre; Ai148-GCaMP6f) (Figure 2A,B). We imaged on average 181+/-77 (mean+/-SD) Slc17a7+ cells or 15+/-10 VIP+ cells per session from a total of 12-13 sessions per transgenic line for each image set (Supplemental Figure 2A). We measured activity in the primary visual cortex, VISp, and one higher visual area, VISal. We did not observe major differences between these two areas for the metrics we evaluated, so data was combined across areas for the analyses reported here. In each imaging session, one of the 4 sets of natural scene images was shown.

**Figure 2:**
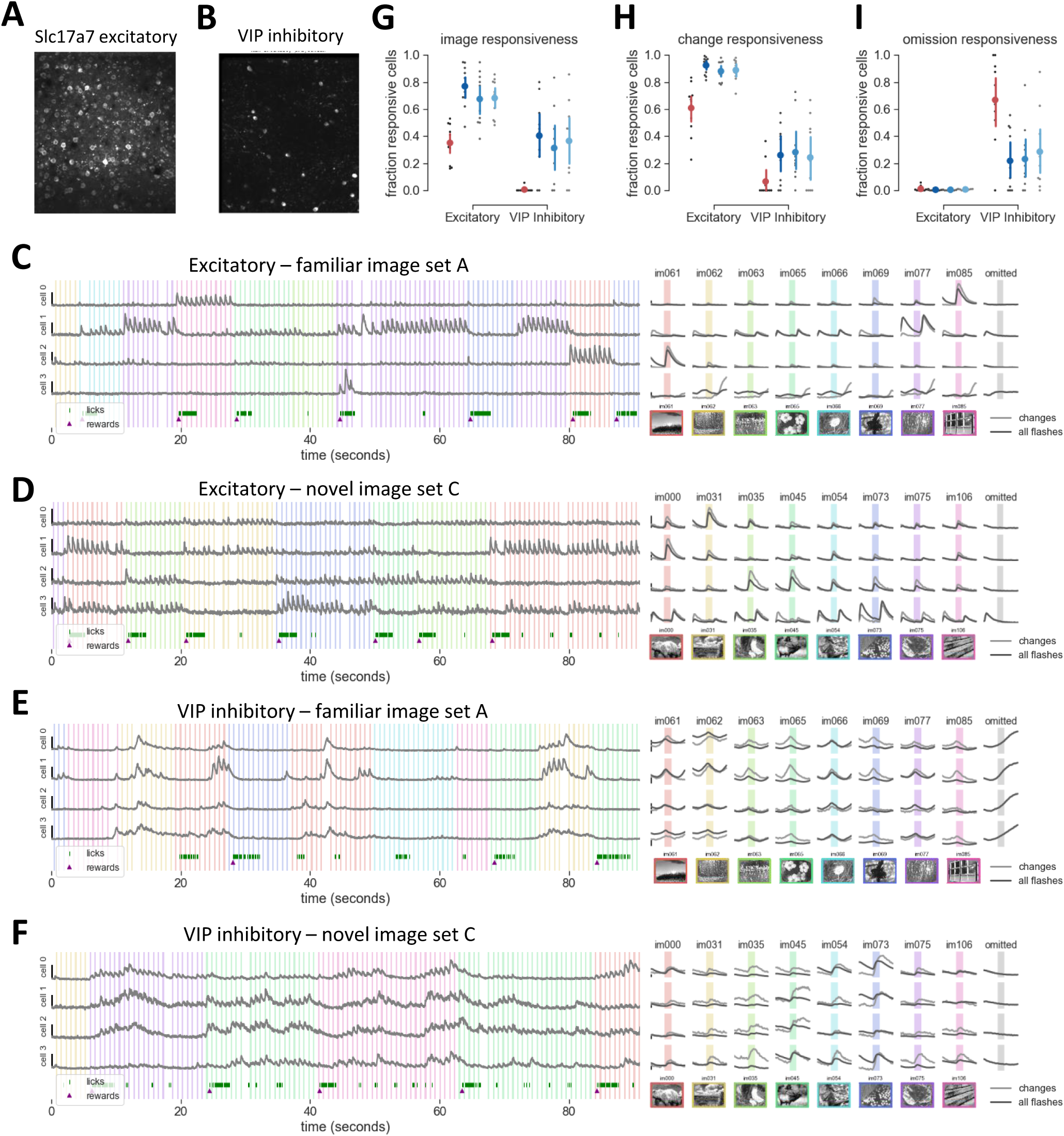
Activity in layer 2/3 excitatory and VIP inhibitory cells during image change detection behavior. **A)** Maximum intensity projection from a 2-photon field of view in layer 2/3 of an Slc17a7- IRES2-Cre;CaMk2-tTa;Ai93 mouse expressing GCaMP6f in excitatory neurons. **B)** Same as A for layer 2/3 inhibitory neurons in a Vip-IRES-Cre; Ai148(GCaMP6f) mouse. **C)** Example excitatory cells from a session with familiar images. Left panel: dF/F traces from 4 excitatory cells over 90 second epoch of a behavior session with the familiar image set A. Scale bars on left indicate 150% dF/F. Colored vertical bars indicate image presentation times; timing of licks and reward delivery are shown at bottom. Right panel: average response of the same 4 cells to each image, as well activity during omission of image (right column, gray indicates time of stimulus omission). Scale bars indicate 25% dF/F. Note the selectivity of excitatory responses and temporal dynamics relative to stimulus onset. **D)** 4 excitatory cells from session with novel image set C. Left panel scale bars indicate 150% dF/F, right panel scale bars indicate 25% dF/F. **E)** 4 VIP cells from a session with familiar image set A. Note the dynamics of the response relative to stimulus onset and stimulus omission. Left panel scale bars indicate 250% dF/F, right panel scale bars indicate 25% dF/F. **F)** 4 VIP cells for a session with novel image set C. Note that stimulus evoked responses are unselective and highly correlated across the population. Left panel scale bars indicate 250% dF/F, right panel scale bars indicate 25% dF/F. **G)** A larger fraction of cells are image responsive for novel image sets compared to the training set A. Responsiveness is defined for each cell as having >25% of preferred image flashes with a significant response compared to a shuffled distribution of stimulus omissions. The fraction of responsive cells is shown for each session in gray, with mean +/- 95% confidence intervals in color. p<0.0005 for all comparisons with image set A for excitatory cells and p<0.01 for all comparisons with image set A for VIP inhibitory cells. **H)** A larger fraction of cells are responsive to the first flash after an image change for novel image sets. Responsiveness computed as described in G, limited to the change flash only. p<0.0006 for all comparisons with image set A for excitatory cells, no significant differences for VIP cells (p>0.05 for all image set comparisons). **I)** Cells with elevated activity during stimulus omission are most common in VIP populations for the trained image set A. Omission responsiveness is defined as having >25% of stimulus omission trials with a significantly larger response than the preceding image flash. p<0.02 for A-B and A-C for VIP cells.

Excitatory cells were typically responsive to only one or a few images in each set (Figure 2C,D). When presented with their preferred stimulus, most excitatory cells responded with an increase in fluorescence shortly after stimulus onset, although some cells responded after stimulus offset. VIP cells were generally less selective for image identity than excitatory cells, and VIP cells were more correlated with each other in their pattern of activity (Figure 2E,F). Moreover, VIP cells showed clear differences in their activity during novel versus familiar image sessions. Stimulus-driven activity was apparent during sessions with novel image sets (Figure 2F) but was reduced or absent with familiar images (Figure 2E). In sessions with familiar image sets, many VIP neurons showed ramp-like responses that preceded stimulus presentation, and these ramping responses were even more pronounced when image flashes were omitted (Figure 2E, right panel). In contrast, during novel image sessions, VIP cells showed little activity when stimuli were omitted (Figure 2F, right panel). Additional examples of image responses from different sessions are shown in Supplemental Figure 2E-H.

Qualitatively, we observed that both layer 2/3 excitatory and VIP inhibitory populations had differences in stimulus responsiveness, image tuning, and neural dynamics during behavioral sessions with novel versus familiar images. These results are further explored and quantified in the sections below.

### Image responses and omission activity during familiar and novel sessions

To quantify image responsiveness for each cell, we compared the mean dF/F value in a 500ms window after each stimulus flash with a shuffled distribution of dF/F values taken from the omitted flash periods when no stimulus was shown. Cells were considered image responsive if they had a significantly larger response for >25% of preferred image flashes. Most excitatory cells were image responsive for novel stimulus sets (Figure 2G; range = 67-77% for image sets B, C, and D), whereas a lower fraction of cells responded to familiar images (35%, image set A). About a third of VIP cells were stimulus responsive for novel images (range = 31-40%), but very few had reliable stimulus-specific activity for familiar images (1%). Note that according to this metric, cells with stronger activity during stimulus omission compared to stimulus presentation will not be considered as image responsive.

We also evaluated change responsiveness for each cell using the same metric, restricting the analysis to the first flashes after a change to each cell’s preferred image (Figure 2H). Cells were considered change responsive if they had a significant response for >25% of change flashes. Most excitatory cells were change responsive during novel image sessions (range = 88-92%, image sets B, C & D), with a smaller fraction for familiar image sessions (61%, image set A). About quarter of VIP cells were responsive for changes to novel images (range = 24-28%), as opposed to only 7% of VIP cells for familiar images.

To quantify activity during omitted flashes we compared dF/F values during the stimulus omission window to the preceding stimulus window. Cells were considered omission responsive if >25% of omission trials had a significantly larger activity compared to the previous image flash (Figure 2I). A large fraction of VIP cells increased their activity following stimulus omission during sessions with the familiar image set (72%, image set A), with a smaller fraction of cells for novel image sets (range = 17-30%, image sets B, C, D). Less than 1.5% of excitatory cells were omission responsive across all image sets.

Thus, we found that novel image sets were associated with increased stimulus responsiveness for both excitatory and VIP cells, while familiar images were associated with greater stimulus omission-related activity in VIP inhibitory cells.

### Mean activity is lower but stimulus selectivity is higher for familiar image sets

Reduced image responsiveness during sessions with familiar versus novel image sets could represent a sparsening of neural representations for the familiar images. Several broad categories of neural sparseness have been described including overall activity, population sparseness, and lifetime sparseness (Willmore et al., 2011). These metrics of sparseness reflect distinct coding properties and do not necessarily co-vary (Willmore and Tolhurst, 2001; Willmore et al., 2011). In the analysis below, we assessed whether each of these metrics varied with image familiarity.

For each cell, we computed the mean response across all presentations of each image and visualized this activity as a heatmap for each image set and cell class (Figure 3A). On average, the strength of evoked activity across the VIP population was ∼4-5 times higher compared to the excitatory population, and excitatory cells were driven by fewer images than VIP cells. We compared the magnitude of responses to familiar and novel image sets within each cell class and found that both excitatory and inhibitory populations had reduced overall activity levels for familiar images (Figure 3B; Supplemental Figure 3A, B).

**Figure 3:**
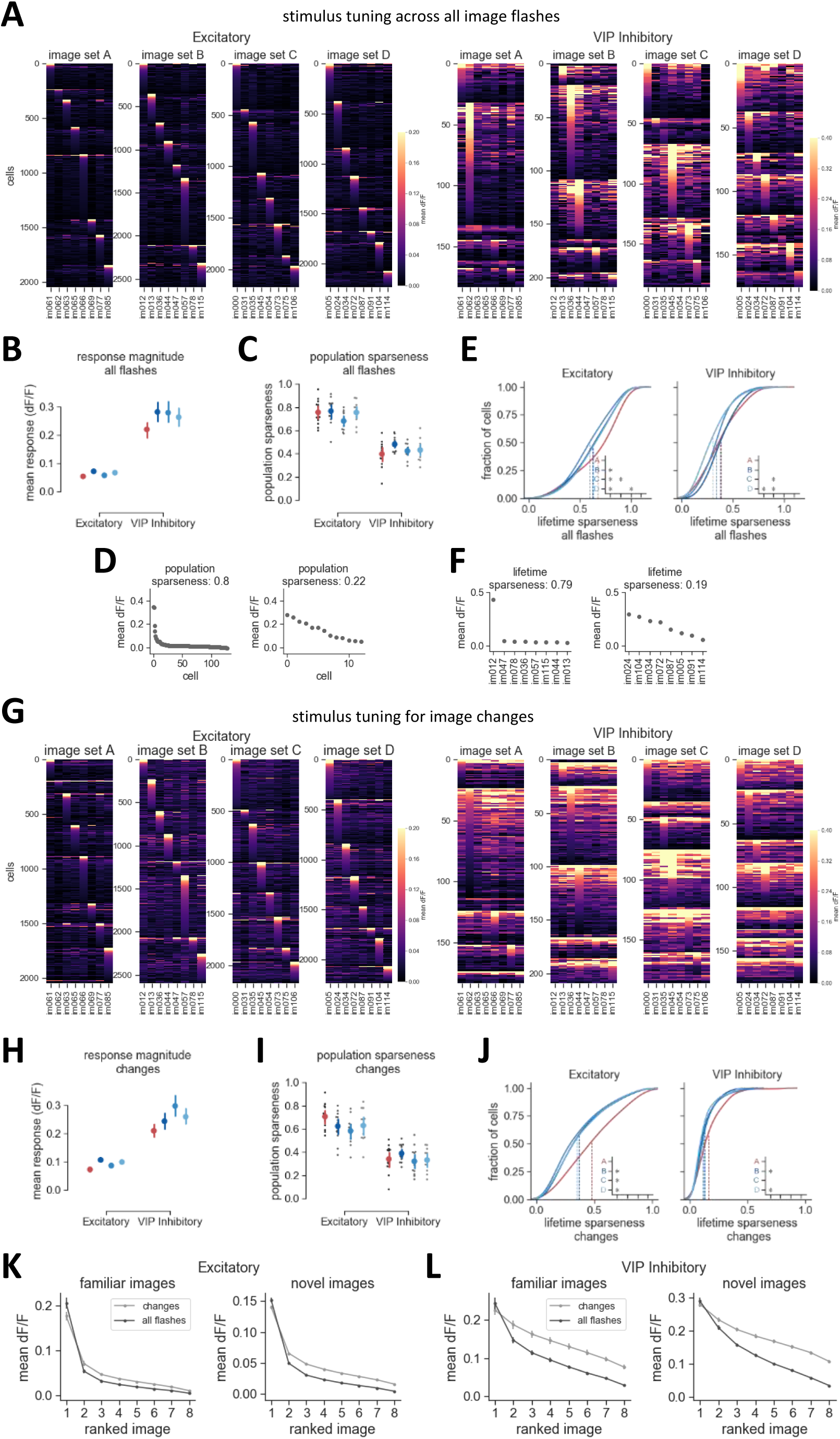
Response sparseness for familiar and novel images. **A)** Cell by image response matrix showing mean response for each image. Response is computed in 500ms window after stimulus onset and averaged over all image flashes. Response for L2/3 excitatory and VIP inhibitory cells are shown for each stimulus set. **B)** Response magnitude across the population is larger for novel image sets. The mean response across all flashes for each cell’s preferred image was computed before averaging across all cells. Error bars show 95% confidence intervals. p<0.001 for comparison of A-B and A-D in excitatory cells, p<0.05 for A-B in VIP cells. **C)** Mean population sparseness across sessions for each cell class and image set. Population sparseness is first computed for each image, then averaged across images within a session. No significant differences are observed across image sets. Gray points are individual sessions, colors indicate mean +/- 95% confidence intervals across sessions. **D)** Example of mean population activity from one session in excitatory cells (left panel) and one session for VIP inhibitory cells (right panel), demonstrating high and low population sparseness values. Each plot shows population activity for a single image. **E)** Cumulative distribution of lifetime sparseness for cells with > 0.05 dF/F for their preferred image. For excitatory cells, inset shows comparisons where p<0.0005. For VIP cells, inset shows where p<0.001. **F)** Example tuning curves from one excitatory cell (left panel) and one VIP inhibitory cell (right panel) demonstrating high and low values of lifetime sparseness. **G)** Same as A) but analysis is restricted to stimulus flashes corresponding to image changes. **H)** Same as B) with analysis restricted to change flashes. p<0.005 for all comparisons with image set A for excitatory cells, p<0.007 for A-C comparison for VIP cells. **I)** Same as C) with analysis restricted to change flashes. No significant differences are observed across image sets. **J)** Same as F) with analysis restricted to change flashes. Inset shows where p<0.001. **K)** Excitatory population tuning curve, averaged across cells with activity > 0.05 dF/F, comparing the image response to changes (light gray) with the average of all flashes (dark gray), for familiar images (left panel) and novel images (right panel) separately. **L)** Same as K) for VIP inhibitory cells.

Next, we computed the average population sparseness value for novel and familiar image sets (Figure 3C). Population sparseness provides a measure of how selectively a population of simultaneously recorded neurons responds to any one stimulus, independently of the overall level of activity (Willmore et al., 2011). Population sparseness of 0 indicates that all neurons respond equally to a given stimulus, whereas a population sparseness value of 1 indicates that only a single cell responds to the stimulus (example sessions and associated population sparseness value are shown in Figure 3D). VIP inhibitory populations had lower population sparseness compared to excitatory populations, as previously reported (de Vries et al., 2018). However, no significant differences were observed across image sets, indicating that experience did not affect population sparseness in these experiments.

To quantify response sparseness of individual cells, we evaluated lifetime sparseness (only active cells with an average dF/F > 0.05 were included in this analysis; Supplemental Figure 3C, D). Lifetime sparseness (Vinje and Gallant, 2000) provides a measure of selectivity on a single neuron basis; at the extremes, this metric takes a value of 0 for cells responding equally to all images and a value of 1 for cells responding to only one image (example cells and associated lifetime sparseness values are shown in Figure 3F). We found that excitatory populations had higher lifetime sparseness values for the familiar image set compared to the novel image sets, and that excitatory cells were typically more sparse than VIP cells (Figure 3E). Plotting the population tuning curve for each image set revealed sharper tuning in excitatory cells for familiar images due to a selective increase in the preferred image response (Supplemental Figure 3E), consistent with previous literature (Woloszyn and Sheinberg, 2012).

To evaluate whether response sparseness differed for image changes, we computed the same metrics but limited analysis to the first image presentation after a change in image identity (Figure 3G). The results paralleled the trends observed for the average of all flashes. The mean activity for image changes was stronger for novel versus familiar image sets (Figure 3H), there was no significant difference in population sparseness across image sets (Figure 3I), and excitatory cells again showed higher lifetime sparseness for familiar images (Figure 3J; Supplemental Figure 3F). However, there were some key differences between image responses averaged across all flashes compared to changes only. Direct comparison of population tuning curves computed from the average of all flashes and population tuning curves for the change flash demonstrate that selectivity for image identity is broader following changes, for both excitatory and VIP cells (Figure 3K,L). This difference was found for both familiar and novel images.

Together, these results demonstrate that while overall population activity levels were reduced for familiar images, single cell selectivity was sharpened. The higher overall level of activity for novel images, combined with lower selectivity, indicates that more cells were recruited to respond to novel images, but the peak response to the preferred image was typically not as high as for images that were experienced during training. Interestingly, population sparseness, which is a measure of the shape as opposed to magnitude of the population activity distribution, was less impacted by experience, indicating that familiar and novel images can be represented by a similar pattern of activity but with a different overall gain. Finally, image tuning was broader for the first flash after a stimulus change compared to the average of all image presentations.

### Enhanced activity following stimulus changes

To explore response dynamics following an image change and subsequent stimulus repetitions, we examined trial averaged activity aligned to the change time for each cell’s preferred image (Figure 4A,B). Many cells showed elevated activity following image changes (see vertical band of activity after time = 0 sec in Figure 4A,B). The response magnitude of excitatory neurons typically decreased on subsequent image flashes, consistent with stimulus specific adaptation, and this was reflected in the population average (Figure 4C). In VIP cells, activity during repetition of the preferred image sometimes increased in magnitude with repeated flashes, and this was reflected in the population average (Figure 4C). However, when averaging over all images, the response to changes was much stronger than subsequent repetitions (Supplemental Figure 4A). The increase in activity with stimulus repetition for preferred images in VIP cells could reflect true facilitation of spiking, or a buildup of calcium activity associated with repetitive stimulation of a strongly driving stimulus. Together, this suggests VIP cells generally have enhanced change responses across many images, but that any change enhancement for the preferred image may be masked by the strong responses to subsequent presentations of the preferred image.

**Figure 4:**
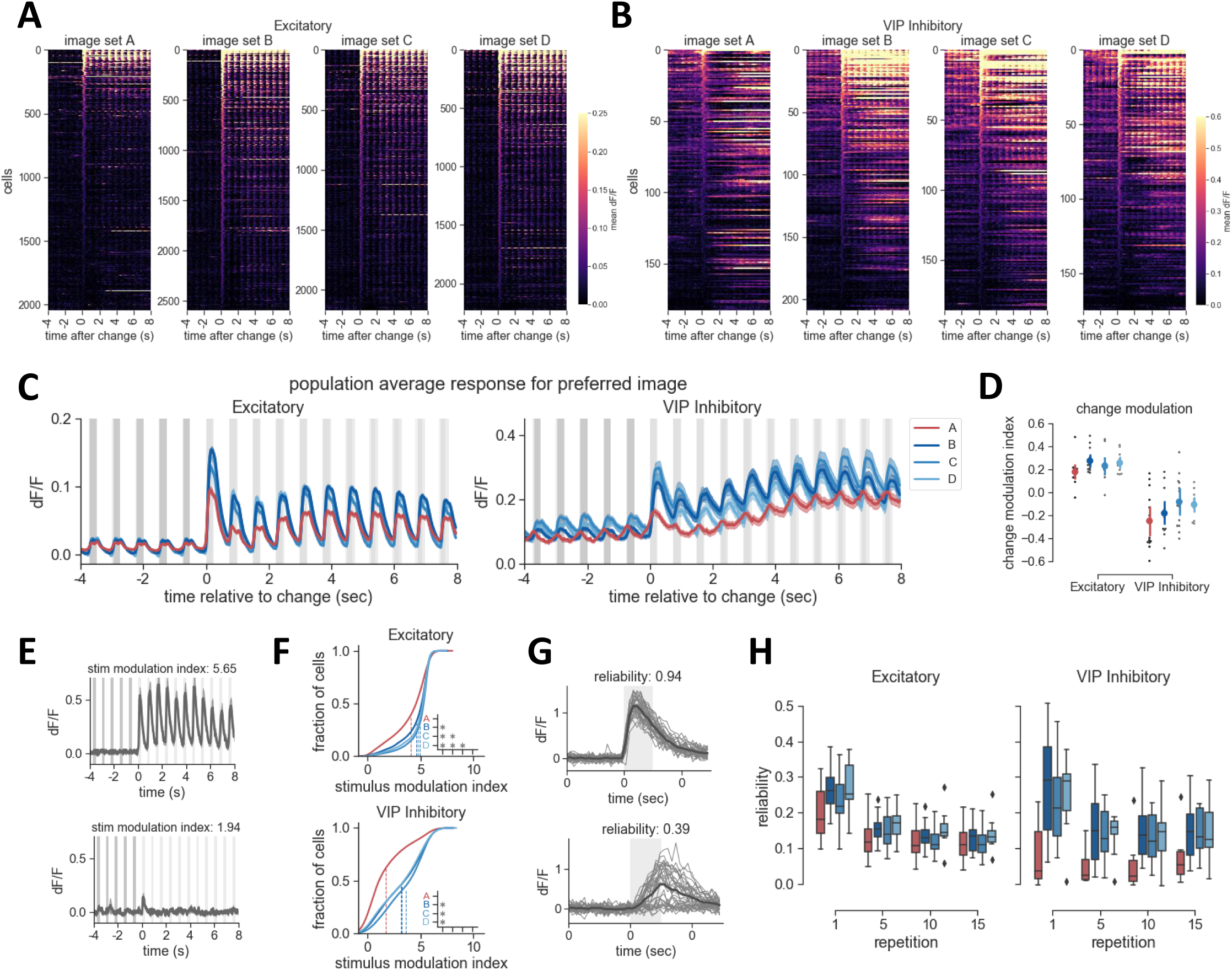
Increased stimulus modulation and reliability for novel images. **A)** Trial averaged response to the preferred image across all excitatory cells, aligned to the time of the change to the preferred image (averaging across multiple images prior to time = 0s). Note the larger fraction of cells with a visible change response for novel image sets B-D. **B)** Trial averaged responses of VIP inhibitory cells for their preferred stimulus, aligned to the change time. Note the increase in stimulus evoked activity for novel image sets B-D. **C)** Population average response for all cells to their preferred image for excitatory and VIP inhibitory populations, aligned to the time of the change to the preferred image. Population response strength is consistently higher for novel image sets compared to the familiar image set A. For excitatory cells, the change response has the largest magnitude followed by a reduction in activity with stimulus repetition. Traces show mean+/-SEM across all cells. **D)** The change modulation index compares the response to the first stimulus following a change with the 10^th^ repetition of that image. The index is computed for the preferred image of each cell, then averaged within a session (gray points). Colored points show mean +/- 95% confidence intervals. No significant differences across image sets for excitatory or VIP. **E)** Example responses demonstrating strong modulation at the stimulus frequency (top panel, modulation index: 5.65 for the 0-8s period after the change time) and weak stimulus modulation (bottom panel, modulation index: 1.94 for t=0-8s). See methods for description of stimulus modulation index. **F)** Cumulative distribution of the stimulus modulation index across cells for each image set by cell class. Responses are more strongly stimulus modulated for novel image sets, especially for VIP populations. Dotted lines indicate the mean of each distribution. Inset shows where p<0.0005. **G)** Example cell responses for the first image flash after a change, showing individual trials for the cell’s preferred stimulus in gray and the trial average in black, demonstrating high reliability (top panel, average trial to trial correlation of the dF/F trace = 0.94), and lower reliability (lower panel, average trial to trial correlation = 0.39). **H)** Responses to novel images and stimulus changes have the highest response reliability. Boxplots show the distribution of mean reliability across sessions, computed first for each cell as a function of stimulus repetition number, then averaged within a session. Repetition 1 is the first flash after an image change and repetition 10 is the 10^th^ image presentation after a change.

To quantify these observations across cells, we computed a change modulation index that compares the response to image changes with the 10^th^ repetition following the change (Figure 4D). When computing this metric for the preferred image for each cell, excitatory populations typically had positive values of this index, whereas VIP cells had negative change modulation index values, reflecting an increasing strength of response with stimulus repetition of the preferred image. However, when considering the mean response across all images, both cell classes had positive values of the index (Supplemental Figure 4B), indicating larger change responses across multiple images. For both changes and the average of all flashes, the strength of the change modulation index was higher for novel image sets in excitatory cells.

These results show that the activity of excitatory and VIP cells is modulated by stimulus repetition following a change, and that enhanced activity for stimulus changes is present across cell classes and image sets. The observed differences in activity following stimulus repetition could reflect stimulus specific adaptation, enhancement associated with a global change signal, or a combination of these factors.

### Activity is more strongly stimulus modulated during sessions with novel images

We noticed that the VIP population response for the familiar image set was only weakly stimulus driven, in comparison to sessions with novel images (Figure 4B,C). To quantify this, we computed a cell by cell metric of stimulus modulation (Matteucci et al., 2019), based on the power at the stimulus frequency (1.33 Hz) over an 8 second window following a change to the preferred image for each cell (Figure 4F). The stimulus modulation index is larger when activity is strongly coupled to the stimulus frequency across repetitive flashes (example cells and associated stimulus modulation index values are shown in Figure 4E). While many VIP cells were strongly stimulus modulated for novel images, modulation at the stimulus frequency was weak for familiar images (Figure 4F, bottom panel), consistent an overall reduction in stimulus driven activity with experience. Excitatory cells also showed reduced stimulus modulation for familiar images, although the effect was not as pronounced as for VIP cells (Figure 4F, top panel).

Trial-to-trial reliability of neural activity can be modulated by bottom-up salience, behavioral context, and learning. Accordingly, we next tested whether response reliability differed for familiar and novel image sets. We quantified reliability for image responsive cells by computing the average pairwise Pearson correlation coefficient across flashes, using the dF/F trace in a 500ms window after stimulus onset (example cells and associated reliability values shown in Figure 4G). Reliability was computed separately for the 1^st^, 5^th^, 10^th^, and 15^th^ flashes after a change to each cell’s preferred image (Figure 4H). We found that reliability was higher for novel image sets in both excitatory and VIP populations, and this result extended across stimulus repetition. Further, reliability was higher for novel images even when only considering the most image selective cells in the population (Supplemental Figure 4C).

Together, these results show that neural activity during task performance with novel, unfamiliar images is stronger across the population, more stimulus driven, and more reliable across trials compared to activity for images that were extensively experienced during training.

### Inter-stimulus activity dynamics of VIP cells are altered by training history

During sessions with familiar images, we observed that VIP cells displayed activity that began ramping up during the gray period prior to image onset (Figure 4C). To further explore these dynamics and their relationship to experience, we examined the average population response of excitatory and VIP cells to all image flashes for each image set (Figure 5A). Excitatory neurons had a sharp stimulus-locked increase in activity following image onset, and although the response magnitude was lower with familiar images, the dynamics were consistent across images sets. In contrast, the dynamics of the VIP population were almost anti-correlated between familiar and novel images sets. With novel images, VIP population activity increased following stimulus onset, but with familiar images, activity increased during the inter-stimulus interval and peaked at stimulus onset. Consistent with this effect, the distribution of peak response times across VIP cells was shifted earlier in time for familiar versus novel images sets (Supplemental Figure 5A).

**Figure 5:**
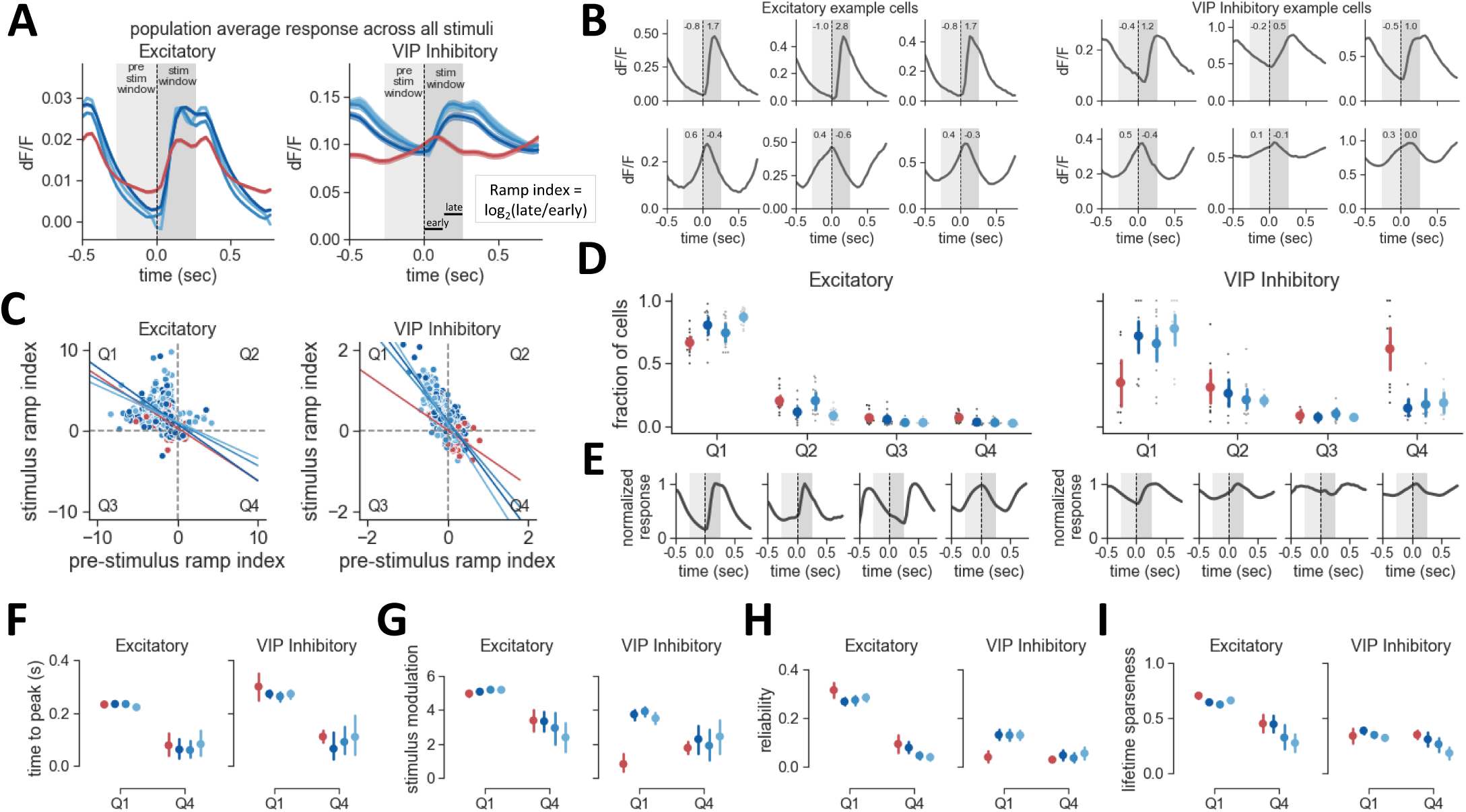
Experience-dependent shift in the dynamics of VIP inhibitory cells. **A)** Population activity averaged over all image flashes for all cells (traces show mean+/-SEM). Note distinct dynamics in VIP population for novel versus familiar images sets. Inset, a ramping index is used to quantify ramping in pre-stimulus and stimulus windows (gray shading); this index compares the dF/F values in the first (early) and second (late) portions of the window. **B)** Example cells showing activity dynamics and associated ramping index values for the pre-stimulus and stimulus periods. Positive values of the ramp index indicate increasing dF/F values in the window, whereas negative values indicate decreasing values. **C)** Relationship of pre-stimulus ramping index (x-axis) with stimulus ramping index (y-axis). Note the negative correlation for both excitatory (r=−0.47 for image set A, r=−0.47 for B, r=−0.42 for C, r=−0.35 for D. p<0.005 for all image sets) and VIP cells (r=−0.61 for image set A, −0.75 for B, −0.66 for C, −0.83 for D. p<0.005 for all image sets). Cells can be separated into 4 quadrants based on pre- and stimulus ramping indices (Q1-4). Only cells with a minimum level of activity (> 0.05 dF/F) in the stimulus window are included in panels C-I. **D)** Fraction of cells belonging to each quadrant across cell class and image set. Cells with pre-stimulus ramping (Q4 cells, increasing activity during pre-stimulus period, negative during stimulus period) are more prevalent for familiar images, particularly for VIP cells, whereas stimulus-evoked responses (Q1 cells, negative ramping during pre-stimulus window, positive during stimulus window) are more prevalent for novel image sets. **E)** Normalized population response for cells belonging to each quadrant from C). **F)** Time to peak is lower for Q4 pre-stimulus ramping cells. **G)** Stimulus modulation index is stronger for Q1 stimulus ramping type cells on average, except for familiar images with VIP. **H)** Trial to trial response reliability across all flashes is higher for stimulus driven Q1 cells compared to pre-stimulus ramping Q4 cells. **I)** Lifetime sparseness for different cell groups. Image selectivity is higher for Q1 cells compared to Q4 cells for excitatory populations, consistent with these cells being a sensory driven population.

Individual cells typically showed either stimulus-evoked responses or ramping activity during the pre-stimulus period that peaked at the time of stimulus onset (Figure 5B). To characterize these dynamics across the population, we made use of a ramping index (Makino and Komiyama, 2015) to quantify activity increases or decreases within the pre-stimulus and stimulus epochs (250ms windows, indicated by light and dark gray shading, respectively, in Figures 5A,B,E). This index compares activity between the early and late portions of a window, giving a positive value for activity increases and a negative value for activity decreases (Figure 5A, right panel inset). While positive pre-stimulus ramping was much less prevalent in excitatory cells than in VIP cells, both cell classes had larger pre-stimulus ramp values and reduced stimulus ramp values for image set A (Supplemental Figure 5B,C), indicating stronger pre-stimulus activity and reduced stimulus activity for familiar images. This ramp index can be unstable for cells with low activity, so all subsequent analysis was limited to active cells that had a mean dF/F greater than 0.05 in the stimulus window for the preferred image.

The values of the ramp index for the pre-stimulus and stimulus windows were typically anti-correlated with each other (Figure 5C; VIP: r = −0.72, −0.78, −0.61, −0.83 for image sets A-D; excitatory: r = −0.29, −0.35, −0.32, −0.26 for image sets A-D; p<0.05 for all comparisons). Cells were naturally divided into four groups based on their pattern of activity before and after stimulus onset (the four groups correspond to the four quadrants of the plot of pre-stimulus versus stimulus ramp index in Figure 5C). For example, cells in quadrant 1 (Q1) had increasing dF/F values in the stimulus window (corresponding to a positive ramp index) and decreasing values for the pre-stimulus window (corresponding to a negative ramp index), consistent with a stimulus-evoked response profile. Cells in quadrant 4 (Q4) were the opposite, having increasing activity prior to stimulus onset and decreasing activity thereafter. The average population response profile across cells belonging to each of these quadrants is shown in Figure 5E.

The majority of excitatory cells for were located in Q1, consistent with strong stimulus-evoked activity (range = 72-84% of cells for image sets A-D; Figure 5D). The majority of VIP inhibitory cells were also in Q1 when tested with novel image sets (range = 69-78% for image sets B-D). However, there was a large increase in the proportion of VIP cells in Q4 showing pre-stimulus ramping for sessions with the familiar image set (55% for image set A compared to 7-9% for image sets B-D; Figure 5D). This represents an experience-dependent shift in inter-stimulus activity of the VIP population. Cells belonging to these distinct response types also differed in their stimulus-evoked response properties. Q4 cells had earlier response times (Figure 5F) and were less image selective (Figure 5I). In contrast, Q1 cells, in line with being sensory-driven, were more stimulus modulated by repeated image flashes, more reliable across trials, and more image selective than Q4 inter-stimulus ramping type cells (Figures 5G-I).

### VIP cells have strong ramping activity during omission of an expected stimulus

Would cells with pre-stimulus ramping continue to ramp if an image was omitted? To assess this, we analyzed activity during periods in which the stimulus flash was randomly omitted. Such trials made up 5% of all stimulus flashes during 2P imaging sessions (with the exception of the change flash and the flash immediately prior to a change).

Strikingly, during stimulus omission in sessions with familiar images, VIP population activity continued to ramp up until the subsequent stimulus flash, more than doubling in response magnitude within the omission window (Figure 6A). This ramping was much stronger during familiar compared to novel image sessions. Omission ramping behavior was not present in the excitatory population on average. While individual excitatory cells could show omission ramping (Figure 6B), these types of responses were extremely rare across the population (Figure 6C). In contrast, for familiar image sessions, most VIP cells showed an increase in activity during the omission period (from time = 0 sec to time =0.75 sec in Figure 6D). In novel image sessions, VIP cell activity was concentrated outside the omission period, with visible stimulus-locked activity in the surrounding timepoints. Still, a subset of VIP cells for image sets B-D showed some degree of activity during the omission period.

**Figure 6:**
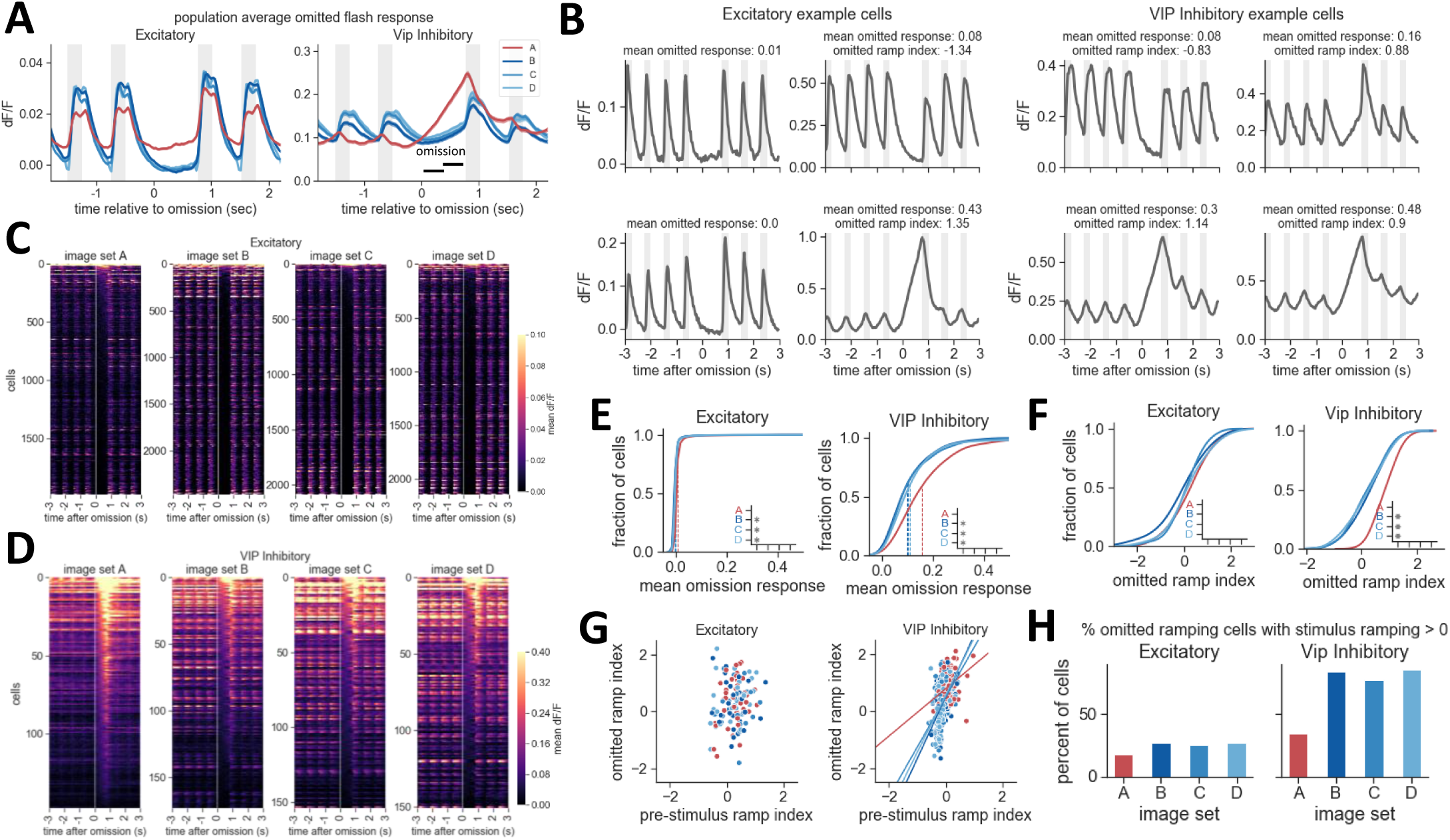
VIP cells show strong ramping activity during stimulus omission. **A)** Average population activity during stimulus omission. On average, excitatory neurons have no change in activity during stimulus omission (left panel). In contrast, activity of the VIP population in sessions with familiar images (image set A) continues to ramp up until the time of the next stimulus presentation. In sessions with novel images, the VIP population also shows little change in activity following stimulus omission (right panel, image sets B-D). Inset, window over which an omission ramp index is computed. Note image-evoked activity before and after the omission for excitatory populations and VIP inhibitory populations for novel images, but not the familiar image set A. **B)** Example cells showing different response dynamics during stimulus (gray bars) and omission periods. Cells typically show either stimulus-evoked activity and no omission response, or pre-stimulus ramping and strong omission responses. Some cells (example on top right) show a combination of stimulus-evoked and omission activity. **C)** Heatmap of activity around the time of stimulus omission across all excitatory cells, sorted by magnitude of activity in the omission window (start of omission period is shown by white vertical line at time = 0 and extends to 750ms thereafter when the next stimulus is presented). Image responses are visible before and after stimulus omission, but activity is near zero for most excitatory cells during stimulus omission. **D)** Heatmap of omission-related activity across all VIP cells. Most VIP cells show ramping during omission for image set A, with fewer cells exhibiting omission ramping in sessions with novel image sets. **E)** Cumulative distribution of mean dF/F activity during the omission window. Dotted lines indicate the mean of each distribution. For excitatory cells, p<0.0005 for all comparisons with image set A. For VIP cells, p<0.002 for all comparisons with image set A. **F)** Cumulative distribution of stimulus omission ramp index showing increased ramping activity for VIP cells with familiar image set A. Note that the omitted ramp index is only computed for cells with a mean dF/F value > 0.05 dF/F during the omission window. No significant differences between image sets for excitatory cells. For VIP cells, p<0.0005 for all comparisons with image set A. **G)** The strength of the omission ramp index (y-axis) and pre-stimulus ramp index (x-axis) are positively correlated, indicating that cells with pre-stimulus activity typically also show ramping during stimulus omission. Fits were not significant for excitatory cells. For VIP, p<0.005 for all image sets, r=0.38 for image set A, r=0.62 for B, r=0.53 for C, r=0.58 for D. **H)** Fraction of omission ramping cells with positive stimulus-evoked activity is higher for novel image sets.

To quantify ramping activity associated with stimulus omission across cells, we computed the ramping index over the stimulus omission window. Here we focused on cells that were active during the omission, with an absolute dF/F of > 0.05 dF/F (3-6% of excitatory cells and 68-76% of VIP cells met this criterion; Figure 6E). Most VIP cells that met the minimum activity level had positive values of the omission ramp index for sessions with familiar images (Figure 6F, right panel). Overall, 80% of VIP cells showed omission ramping for familiar sessions, compared to 40.5% of cells for novel sessions on average. While ramping was rare in excitatory cells, the fraction of cells with ramping during omission was slightly increased for sessions with familiar images (1.1% for familiar, 0.6% for novel on average).

When looking at the response profiles of individual cells (Figure 6B), we noticed a relationship between omission ramping and stimulus-related dynamics. Typically, cells that had strong stimulus-evoked responses showed little to no activity in the omitted window, whereas cells with pre-stimulus activity had the strongest ramping during stimulus omission (Figure 6B). Indeed, there was a correlation between the pre-stimulus ramping index and the omission ramping index for VIP cells (Figure 6G). However, the slope of this relationship differed between familiar and novel images. To explore this difference, we computed the fraction of cells that had both a positive omission ramp index and a positive stimulus ramp index (Figure 6H) and found that the majority of VIP cells that show omission ramping during novel image sessions also have increasing activity following stimulus presentation (example cell showing this profile in upper right panel of Figure 6B). This indicates that during visual stimulation with novel images, VIP cells are more likely to show a combination of sensory driven activity and ramping during omission.

Overall, these results demonstrate that the dynamics of VIP cells in visual cortex are strongly impacted by experience.

## DISCUSSION

By imaging cortical activity in L2/3 excitatory and VIP inhibitory neurons during a visual task with familiar and novel images, we identified several changes associated with training history. Extended behavioral experience with natural scene images resulted in reduced overall activity levels and increased selectivity of single neuron representations. In contrast, responses to novel images were strong and reliable, particularly for stimulus changes. Interestingly, VIP cells had distinct activity dynamics and stimulus omission responses when tested with familiar versus novel images. Novel images drove stimulus-locked activity in VIP cells, whereas VIP cells were primarily active during the inter-stimulus interval during sessions with familiar images. Together these results indicate that both stimulus coding and temporal dynamics of visual cortical circuits can be impacted by experience in a cell type-specific manner.

### Distinct activity and tuning in excitatory versus VIP cells

Across all conditions, VIP cells had higher overall average activity levels compared to excitatory neurons. In contrast, excitatory neurons were more strongly stimulus modulated and had more reliable stimulus-evoked responses. Excitatory cells were sharply tuned for natural images, whereas VIP cells responded more broadly, and were more correlated with each other. These results are consistent with previous studies showing overall activity and selectivity differences of VIP compared to excitatory cells (Kerlin et al., 2010; de Vries et al., 2018). We also observed that stimulus repetition influenced activity: both cell classes showed stronger responses to changes relative to subsequent stimulus repetitions, and the change response was more broadly tuned than the adapted response (10th stimulus repetition). These differences could be a result of stimulus specific adaptation (Grill-Spector et al., 2006; Nelken and Ulanovsky, 2007), or novelty enhancement (Hamm and Yuste, 2016; Homann et al., 2017; Vinken et al., 2017), potentially reflecting a combination of bottom-up, top-down, or neuromodulatory influences (Ranganath and Rainer, 2003).

### Experience reduces overall activity and sharpens tuning

Whereas stimulus repetition was associated with reduced activity on short timescales (∼seconds), we also observed reduced activity in both VIP and excitatory cells due to long-term experience with highly familiar images (∼days-weeks). The fraction of image responsive cells (Figure 2) and response magnitude across the population (Figure 3, Figure 4) was lower with familiar versus novel images. Previous studies have shown reductions in activity with experience (Anderson et al., 2008; Mruczek and Sheinberg, 2007; Woloszyn and Sheinberg, 2012), and enhancement for novelty (Hamm and Yuste, 2016; Homann et al., 2017; Ranganath and Rainer, 2003). A recent study found that experience-induced reductions in the fraction of active cells can be specific to distinct functional subpopulations: ‘transient’ cells were reduced with experience, but ‘sustained’ and ‘suppressed by contrast’ cells showed no change in the fraction of active cells (Weskelblatt and Niell, 2019). Reduced activity for highly familiar stimuli may serve to more efficiently code for stimuli which are predictable, utilizing a smaller population of cells to represent learned information (LeMessurier and Feldman, 2018). On the other hand, enhanced activity for novel stimuli may be involved in the detection of salient and behaviorally meaningful events by augmenting output to downstream targets and facilitating associative plasticity (Ranganath and Rainer, 2003). Consistent with this idea, we observe a nearly two-fold increase in the number of cells that respond to novel images compared to familiar ones, for both excitatory and VIP inhibitory populations (Figure 2G,H).

We found that stimulus selectivity of single cells, measured by lifetime sparseness, was higher in L2/3 excitatory cells when tested with familiar images. Stimulus experience and task learning have been shown to increase orientation selectivity in the visual cortex of mice (Fiser et al., 2016; Frenkel et al., 2006; Jurjut et al., 2017; Khan et al., 2018; Poort et al., 2015) and selectivity for natural images including complex objects in primates (Freedman et al., 2006; Ghose et al., 2002; Meyer et al., 2014; Schoups et al., 2001; Yang and Maunsell, 2004). Although few studies have explored cell type-specific differences in stimulus tuning following experience, one study in macaque IT showed that both regular spiking (putative excitatory) and fast spiking (putative inhibitory) cells showed increased stimulus selectivity for familiar images (Woloszyn and Sheinberg, 2012). A study in mice reported increased selectivity with experience in excitatory and multiple inhibitory cell types, including PV and SST, but did not find a change in selectivity for VIP cells (Khan et al., 2018).

Many studies have shown that experience-dependent changes in responsiveness and selectivity can underlie improvements in perception and behavior. Here, we did not observe significant differences in behavioral performance between familiar and novel image sets. Mice rapidly generalized task performance to novel images and had comparable hit and false alarm rates, reaction times, and patterns of running behavior across image sets (Figure 1, Supplemental Figure 1). Accordingly, it is unlikely that the differences in response properties for familiar versus novel images are the consequence of differences in animal behavior or task performance. Consistent task performance despite differences in stimulus coding suggests increased activity for novel images could help maintain task performance in a new stimulus context, or alternatively, that a more efficient representation is sufficient for performance with familiar images.

### Experience alters temporal dynamics of VIP cells

One of the main findings in this study is that extended visual experience with natural scene stimuli leads to altered activity dynamics in VIP inhibitory cells in layer 2/3 of visual cortex (Figure 5). VIP populations were largely stimulus-driven by novel images but showed prominent inter-stimulus ramping activity with familiar images. This represents a major experience-induced change in response dynamics that has not been demonstrated previously. We found an inverse relationship between inter-stimulus ramping and stimulus-triggered ramping in VIP cells, indicating a trade-off between sensory-driven and pre-stimulus activity (Figure 5). Further, cells with inter-stimulus ramping continue to increase their activity following stimulus omission, up until the time of the next stimulus onset (Figure 6). The magnitude of omission-related activity was several times larger than stimulus-driven activity, suggesting these are meaningful signals that could strongly influence network activity.

What does pre-stimulus and stimulus-omission ramping activity in VIP cells represent? One possibility is that this activity reflects the temporal structure of the behavioral task such that these signals encode predictions about stimulus timing or reward expectation, or serve as a general attentional signal (Nobre and Van Ede, 2018). Previous studies have described stimulus and reward expectation signals in the visual cortex of rodents, including an early example showing that pairing visual stimulation with a temporally predictable reward produces reward timing signals in visual cortex (Shuler and Bear, 2006). Cholinergic signaling has been implicated in mediating these changes (Chubykin et al., 2013), as well as other attention and learning dependent effects (Hasselmo, 1995; Lee and Dan, 2012). Given the strong neuromodulatory drive to VIP cells, including cholinergic input, neuromodulation is a candidate mechanism to underlie the shift in response dynamics we observe across the VIP population. Visual cortex has also been shown to learn experience-dependent stimulus predictions for repeated sequences of visual stimuli (Gavornik and Bear, 2014; Xu et al., 2012). However, in these studies, the predictive signal peaked at the expected time of the predicted event on omission trials, whereas our results show a continued ramping past the expected time of stimulus onset on omission trials. This suggests that the pre-stimulus ramping and associated omission ramping signals we observe may represent something other than a pure prediction of timing of stimulus or reward.

In virtual navigation paradigms in which mice locomote along a linear track, V1 neurons have been found to predict upcoming stimuli at specific locations, with separate populations of cells that respond when expected stimuli are omitted (Fiser et al., 2016). In contrast, we observe a correlation between pre-stimulus and omission related activity on a cell by cell basis, specifically for VIP cells (Figure 6G), and these cells are generally unselective for image identity (Figure 5I). Another study using virtual navigation with a visual discrimination task found pre-stimulus ramping activity specifically in the subpopulation of excitatory cells that encoded the rewarded stimulus, suggestive of reward anticipation (Poort et al., 2015). The emergence of an experience dependent ramping profile in anticipation of an imminent foot shock has also been documented (Makino and Komiyama, 2015). However, without an omission condition, it is unclear how these cells would behave if the expectation was violated. Further modifications of our change detection task to include omitted rewards, or to vary the inter-stimulus interval, could help to distinguish between coding of stimulus timing versus reward anticipation. Another open question is whether pre-stimulus ramping behavior emerges following passive exposure to the same set of familiar images in the absence of reward, or whether active task performance and reinforcement are necessary for the emergence of this phenomenon.

It is important to note that most prior studies documenting predictive or ramping activity reflect measurements from excitatory neurons, and thus may not be directly comparable to our results in VIP cells. In fact, it is surprising that we do not observe many excitatory cells with pre-stimulus or omission related activity given these prior results. One study of inhibitory cells in visual cortex observed that VIP population activity peaked near stimulus onset in an orientation discrimination task, similar to our results for familiar images, but did not note a major change in VIP activity dynamics before and after learning (Khan et al., 2018). In the prefrontal cortex of the mouse, VIP cell activity during the delay period of an auditory go/no-go discrimination task was found to be important for behavior, and activation of VIP cells during the delay improved task performance by improving coding in the excitatory population (Kamigaki and Dan, 2017). The inter-stimulus activity of VIP cells during sessions with familiar images in the change detection task may be similarly important for enhancing the responses of subsets of excitatory cells to an expected change stimulus, via disinhibition through SST cells. In contrast, during sessions with novel images, VIP activity is driven by stimulus presentation and could serve to increase the gain of stimulus evoked excitatory responses to facilitate learning of new images via associative plasticity (LeMessurier and Feldman, 2018; Ranganath and Rainer, 2003). Future studies examining the evolution of VIP activity across multiple behavior sessions as novel images become familiar, as well as concurrent recordings of VIP and excitatory cells and other inhibitory classes including SST cells, will be critical to determine the time course and mechanistic nature of these interactions.

### Predictive coding and experience-dependent changes in activity

Predictive processing has emerged as a powerful paradigm for understanding brain function and may help reconcile the traditional view of sensory processing with increasing evidence for experience and context dependent modulation in early sensory areas. This family of theories posits that the brain constructs an internal model of the environment based on experience, and that incoming sensory information is compared with learned expectations to continually update the model (De Lange et al., 2018; Keller and Mrsic-Flogel, 2018; Lochmann and Deneve, 2011; Rao and Ballard, 1999). This dynamic updating with experience may be associated with a shift in the balance of bottom-up sensory and top-down predictive pathways as internal representations become more effective at predicting the external causes of sensations. As stimuli become familiar with learning, predictive signals are thought to ‘explain away’ incoming information by suppressing bottom-up input, resulting in a sparse code. In contrast, novel or surprising stimuli are expected to robustly drive neural activity, signaling deviations from learned predictions. Our results are consistent with this model, demonstrating reduced activity with long-term experience, in addition to within-trial stimulus specific adaptation. Further, the observation of a switch between stimulus evoked activity and inter-stimulus ramping in VIP cells provides a novel demonstration of an experience dependent change in activity dynamics in a specific subtype of inhibitory interneuron that may regulate the balance between top-down predictive and bottom-up sensory information. Future studies examining the activity and impact of the diverse inputs to VIP cells, including neuromodulatory inputs from subcortical structures and feedback projections from other cortical regions, will be important to establish the function of and mechanism behind the shift in VIP dynamics with experience.

## FIGURE LEGENDS

**Supplemental Figure 1:**
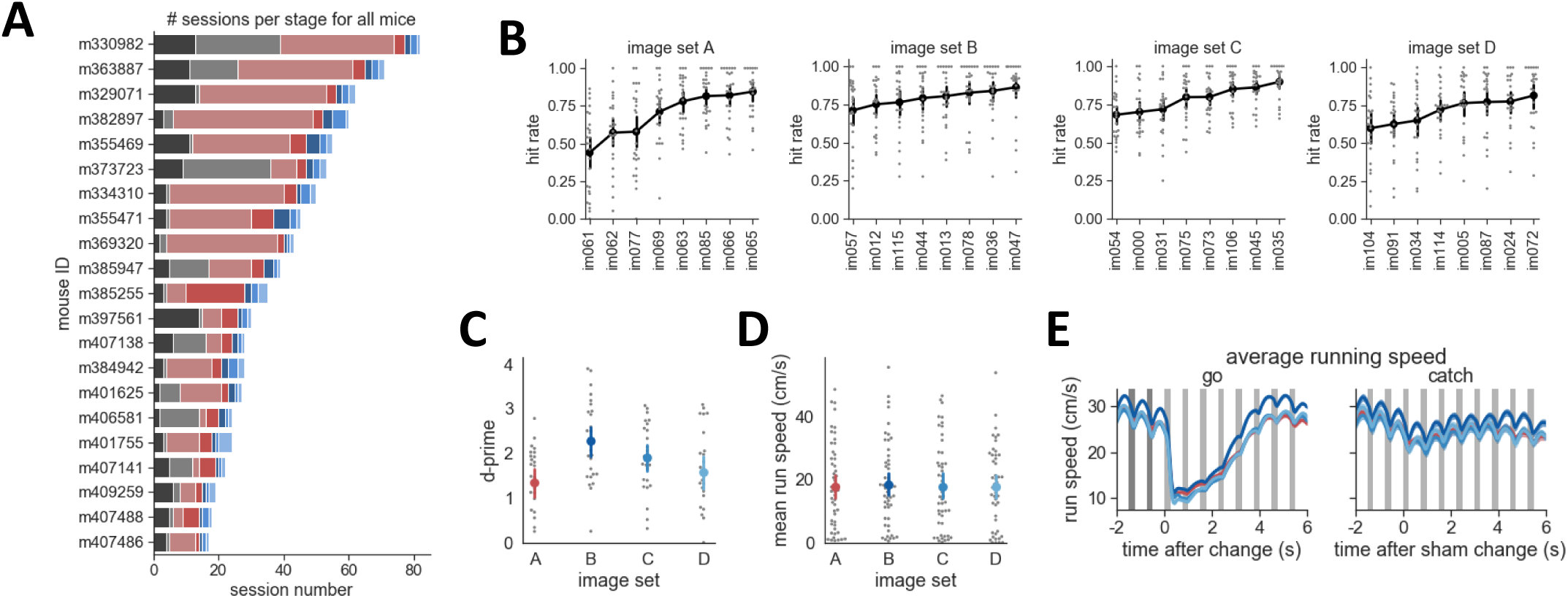
Behavior performance across image sets. **A**) Number of sessions in each training stage for all mice in this study. **B)** Hit rate for each of the 8 images in the 4 image sets. Mean and 95% confidence intervals are shown in black with gray points for individual behavior sessions. **C)** Mean d-prime is similar across image sets (no significant difference across image set pairs except A-B, where p=0.002). **D)** Mean running speed is similar across image sets (p>0.05 for all comparisons). Error bars in B-D are 95% confidence intervals. **E)** Task-related running behavior is similar across image sets. Average running trajectory relative to a stimulus change is plotted for each image set. Average +/- SEM for go and catch trials (n=6260+/-294 go trials, n=941+/-40 catch trials).

**Supplemental Figure 2:**
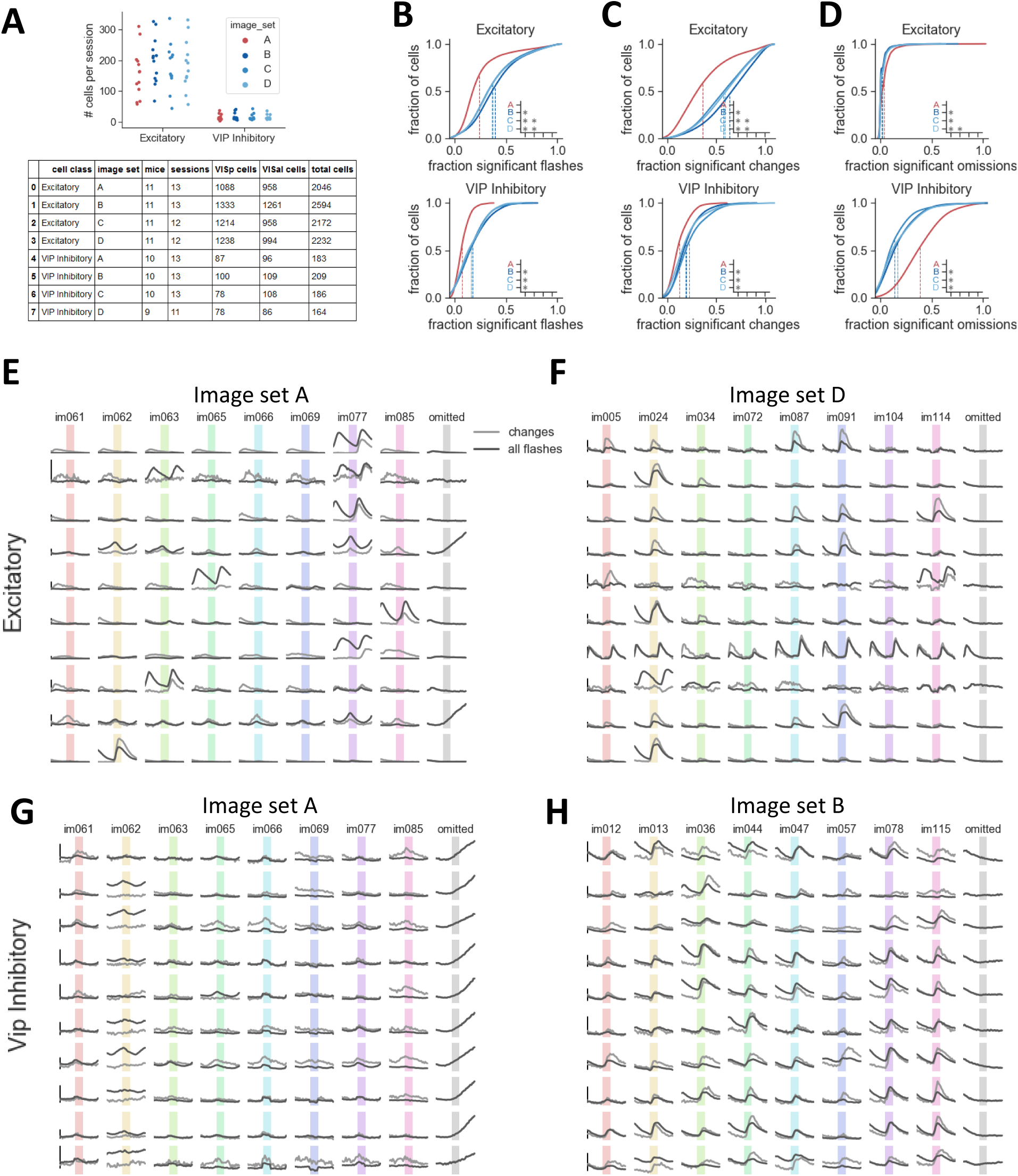
Image, change, and omission responsiveness. **A)** Top: number of segmented ROIs per imaging session by cell class and image set. Bottom: table of experimental sessions including image set, cell class, and area. **B)** Distribution of significant responses to flashes of each cell’s preferred image. Dotted lines indicate the mean. Inset shows image set comparisons where p<0.0002. **C)** Distribution of significant change trials across cells. Inset shows image set comparisons where p<0.0002. **D)** Distribution omission responsive trials across cells. Very few excitatory cells respond during stimulus omission, while a large number of VIP cells show omission related activity, especially for the familiar image set. **E)** 10 excitatory cells from one session with familiar image set A. Average response across all presentations of each image (columns, colored bars indicate stimulus presentation) and during stimulus omission (right column, gray bar indicates time of stimulus omission). Light gray shows response for image change flash for each image in light gray. Cells with highest SNR values were selected for display, where SNR is the ratio of mean and standard deviation of dF/F trace. **F)** 10 excitatory cells from a session with novel image set D. **G)** 10 VIP inhibitory cells from a session with familiar image set A. **H)** 10 VIP inhibitory cells from a session with novel image set B. Scalebars in E-H indicate 25% dF/F.

**Supplemental Figure 3:**
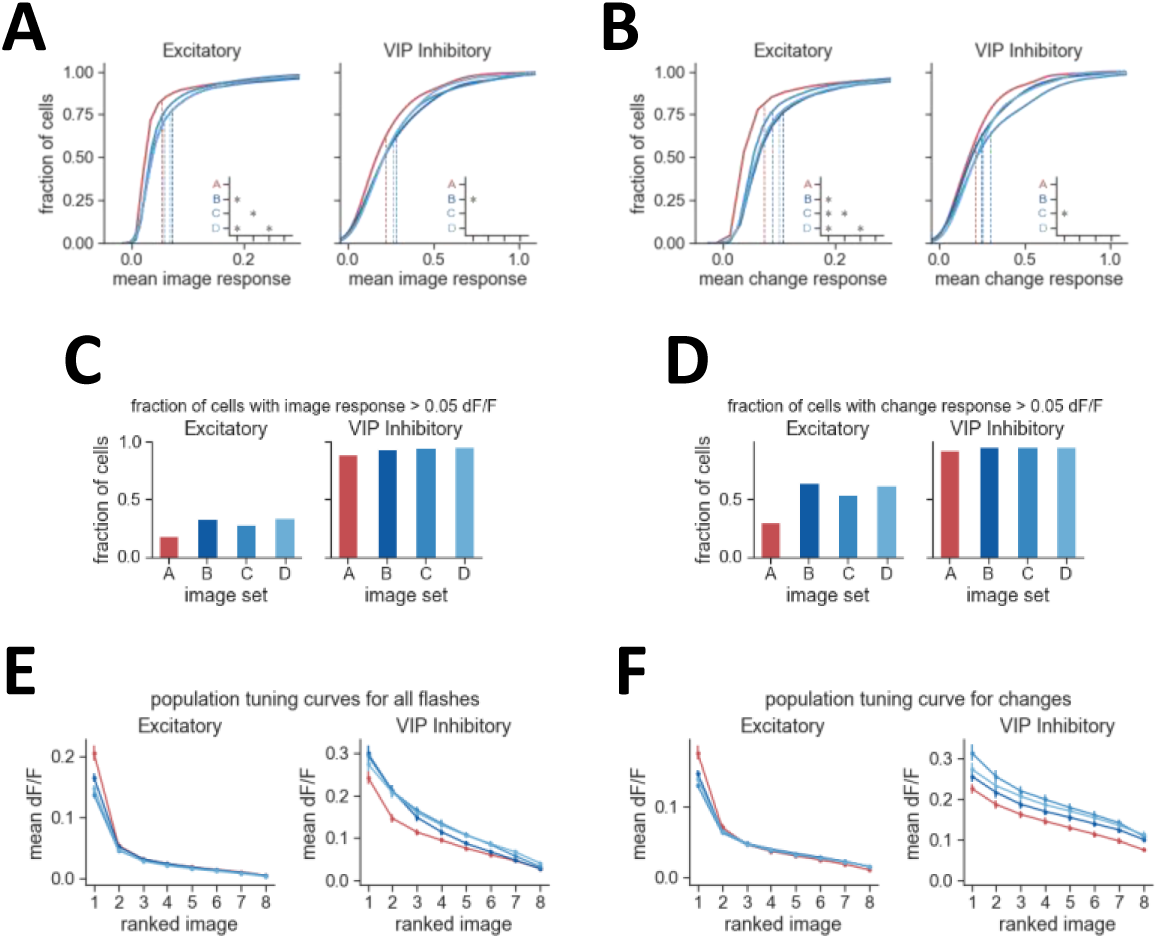
Response strength for familiar and novel images. **A)** Distribution of image response magnitude across cells, taken as the mean dF/F value over all flashes for each cell’s preferred image. Inset indicates where p<0.05. **B)** Distribution of change response magnitude, taken as the mean dF/F value over all change flashes for each cell’s preferred image. **C)** Fraction of cells with a mean image response > 0.05 dF/F for their preferred image. **D)** Fraction of cells with a mean change fesponse > 0.05 dF/F for their preferred image. **E)** Population tuning curve across image sets, averaged across all flashes for cells with > 0.05 dF/F image response**. F)** Population tuning curve across image sets, averaged across change flashes for cells with > 0.05 dF/F change response.

**Supplemental Figure 4:**
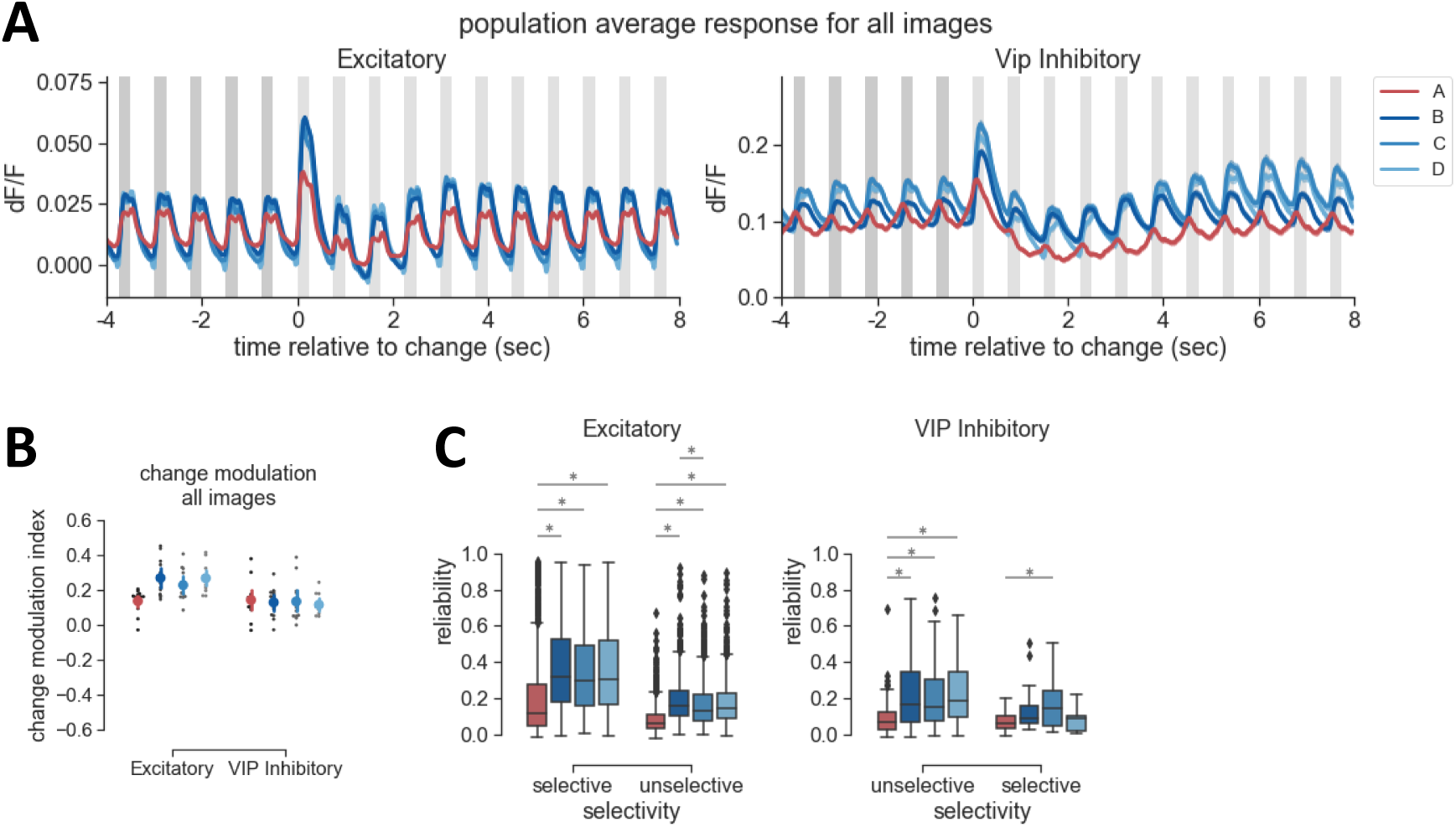
Change modulation across images. **A)** Population average response for all cells across all images for excitatory and VIP inhibitory populations, aligned to the change time. Population response strength is consistently higher for novel image sets compared to the familiar image set A. For excitatory cells, the change response has the largest magnitude followed by a reduction in activity with stimulus repetition. Traces show mean+/-SEM across all cells. **B)** Change modulation index comparing the 1^st^ flash with the 10^th^ flash across all images, averaged across cells within each session. Colored points show mean +/- 95% confidence intervals, gray points show individual sessions. No significant differences across image sets for excitatory or VIP. p<0.05 for comparison of A-B and A-C for excitatory cells. **C)** Responses to novel images (blue box plots) show higher trial to trial reliability compared to familiar images (red blox plots), for both image selective (lifetime sparseness > 0.3) and unselective (lifetime sparseness < 0.3) populations. Boxplots show distribution across responsive cells for each condition. Asterix on plot indicates comparisons where p<0.05.

**Supplemental Figure 5:**
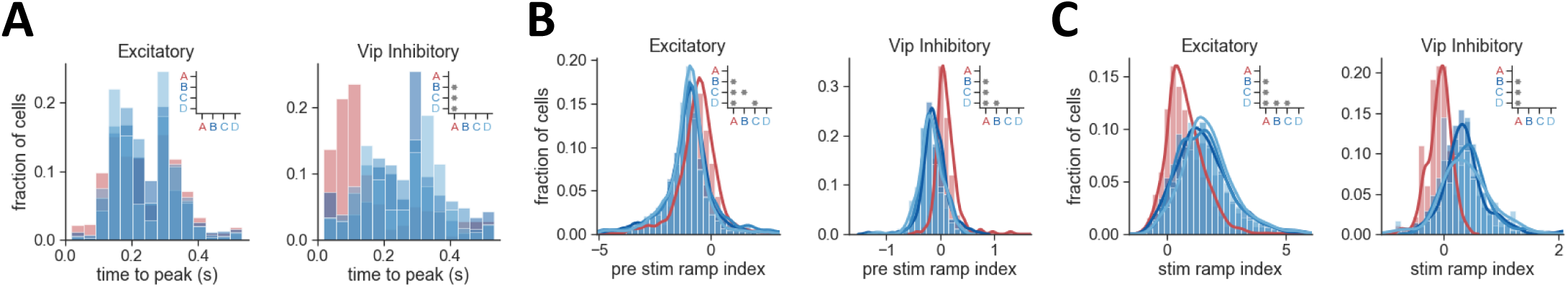
Activity dynamics depend on experience. **A)** Distribution of time to peak response across cells is shifted towards the right for familiar images (red) across the VIP population. Note the bi-modality of response timing for excitatory cells, likely corresponding to on- and off-responsive populations. No significant differences are observed across image sets for excitatory cells. For VIP cells, p<0.0005 for all comparisons with image set A. **B)** Distribution of pre-stimulus ramp index is shifted towards more positive values in sessions with image set A for both excitatory and VIP inhibitory populations, indicating an increased prevalence of predictive signals. For excitatory cells, p<0.0005 for all comparisons across image sets except B-D. For VIP cells, p<0.0005 for all comparisons with image set A, p<0.01 for B-D. **C)** Distribution of stimulus ramp index is shifted towards more positive values for novel image sets for both excitatory and VIP inhibitory populations, consistent with increased stimulus responsiveness for novel images. For excitatory cells, p<0.001 for all comparisons across image sets except B-D. For VIP cells, p<0.0005 for all comparisons with image set A.

## ACKNOWLEDGEMENTS

We thank Jerome Lecoq and Kevin Takasaki for technical help with the 2-photon microscope, Derric Williams for help with behavior and stimulus control software, Douglas Kim, Janelia Research Campus, Howard Hughes Medical Institute, for providing GCaMP6f, and Saskia de Vries, Brian Hu, and Christof Koch for comments on the manuscript. The authors thank the Allen Institute founder, Paul G. Allen, for his vision, encouragement, and support

## MATERIALS AND METHODS

### Mice

All experiments and procedures were performed in accordance with protocols approved by the Allen Institute Animal Care and Use Committee. We used male and female transgenic mice expressing GCaMP6f in VIP inhibitory interneurons (double transgenic: VIP-IRES-Cre x Ai148 mice; https://www.jax.org/strain/010908; https://www.jax.org/strain/030328) or in excitatory glutamatergic neurons (triple transgenic: Slc17a7-IRES2-Cre x CaMKII-tTA x Ai93; https://www.jax.org/strain/023527, https://www.jax.org/strain/010712, https://www.jax.org/strain/024108) were used in these experiments. Mice were single housed and maintained on a reverse 12-hour light cycle (off at 9am, on at 9pm) and all experiments were performed during the dark cycle.

### Surgery

Surgical procedures were performed as described in (de Vries et al., 2018). Headpost and cranial window surgery was performed on healthy mice that ranged in age from 5-12 weeks. Preoperative injections of dexamethasone (3.2 mg/kg, S.C.) were administered at 12h and 3h before surgery. Mice were initially anesthetized with 5% isoflurane (1-3 min) and placed in a stereotaxic frame (Model# 1900, Kopf, Tujunga, CA), and isoflurane levels were maintained at 1.5-2.5% for surgery. An incision was made to remove skin, and the exposed skull was levelled with respect to pitch (bregma-lambda level), roll and yaw. The stereotax was zeroed at lambda using a custom headframe holder equipped with stylus affixed to a clamp-plate. The stylus was then replaced with the headframe to center the headframe well at 2.8 mm lateral and 1.3 mm anterior to lambda. The headframe was affixed to the skull with white Metabond and once dried, the mouse was placed in a custom clamp to position the skull at a rotated angle of 23° such that visual cortex was horizontal to facilitate the craniotomy. A circular piece of skull 5 mm in diameter was removed, and a durotomy was performed. A coverslip stack (two 5 mm and one 7 mm glass coverslip adhered together) was cemented in place with Vetbond (Goldey et al., 2014). Metabond cement was applied around the cranial window inside the well to secure the glass window. Post-surgical brain health was documented using a custom photo-documentation system and at one, two, and seven days following surgery, animals were assessed for overall health (bright, alert and responsive), cranial window clarity, and brain health. After 1-2 week recovery from surgery animals underwent intrinsic signal imaging for retinotopic mapping, then entered into behavioral training.

### Intrinsic signal imaging

Intrinsic signal imaging (ISI) was performed as described in (de Vries et al., 2018) to produce a retinotopic map to define visual area boundaries and target *in vivo* two-photon calcium imaging experiments to the center of visual space in each imaged area. Mice were lightly anesthetized with 1-1.4% isoflurane administered with a somnosuite (model #715; Kent Scientific, CON). Vital signs were monitored with a Physiosuite (model # PS-MSTAT-RT; Kent Scientific). Eye drops (Lacri-Lube Lubricant Eye Ointment; Refresh) were applied to maintain hydration and clarity of eye during anesthesia. Mice were headfixed for imaging.

The brain surface was illuminated with two independent LED lights: green (peak λ=527nm; FWHM=50nm; Cree Inc., C503B-GCN-CY0C0791) and red (peak λ=635nm and FWHM of 20nm; Avago Technologies, HLMP-EG08-Y2000) mounted on the optical lens. A pair of Nikon lenses lens (Nikon Nikkor 105mm f/2.8, Nikon Nikkor 35mm f/1.4), provided 3.0x magnification (M=105/35) onto an Andor Zyla 5.5 10tap sCMOS camera. A bandpass filter (Semrock; FF01-630/92nm) was used to only record reflected red light onto the brain.

A 24” monitor was positioned 10 cm from the right eye. The monitor was rotated 30° relative to the animal’s dorsoventral axis and tilted 70° off the horizon to ensure that the stimulus was perpendicular to the optic axis of the eye (Oommen and Stahl, 2008). The visual stimulus for mapping retinopy was a 20° x 155° drifting bar containing a checkerboard pattern, with individual square sizes measuring 25°, that alternated black and white as it moved across a mean-luminance gray background. The bar moved in each of the four cardinal directions 10 times. The stimulus was warped spatially so that a spherical representation could be displayed on a flat monitor (Marshel et al., 2011).

After defocusing from the surface vasculature (between 500 μm and 1500 μm along the optical axis), up to 10 independent ISI timeseries were acquired and used to measure the hemodynamic response to the visual stimulus. Averaged sign maps were produced from a minimum of 3 timeseries images for a combined minimum of 30 stimulus sweeps in each direction (Garrett et al., 2014).

The resulting ISI maps were automatically segmented by comparing the sign, location, size, and spatial relationships of the segmented areas against those compiled in an ISI-derived atlas of visual areas. A cost function, defined by the discrepancy between the properties of the matched areas, was minimized to identify the best match between visual areas in the experimental sign map and those in the atlas, resulting in an auto-segmented and annotated map for each experiment. Manual correction and editing of the results included merging and splitting of segmented and annotated areas to correct errors. Finally, target maps were created to guide *in vivo* two-photon imaging location using the retinotopic map. The center of retinotopic space was computed from azimuth and altitude maps and adjusted for variability in eye position relative to the monitor by zeroing to the anatomical center V1. The corresponding retinotopic location was identified for each visual area, and two-photon imaging was targeted to a region within 20° of the center of gaze.

### Behavior Training

#### Water restriction and habituation

Throughout behavior training mice were water-restricted in order to maintain consistent motivation to learn and perform the behavioral task (Guo et al., 2014). Prior to water restriction mice were weighed once daily for three days to obtain a stable, initial baseline weight. During the first week of water restriction mice were handled daily and habituated to increasing duration of head fixation in the behavior enclosure over a five-day period. Thus, the first day of behavior training occurred after 10 days of water restriction. Mice were trained 5 days per week and could earn as much water as possible during the daily one hour sessions; supplemental water was provided if earned volume fell below 1.0 mL and/or body weight fell under 80-85% of their initial baseline weight. On non-training days mice were weighed and received enough water provision to reach their target weight of 80-85% (never less than 1.0 mL per day).

#### Apparatus

Headposted mice were trained in custom-designed, sound-attenuating behavior enclosures. Visual stimuli were displayed on a 24” LCD monitor (ASUS, Model # PA248Q) placed at a ∼15cm distance from the mouse’s right eye. The monitor was rotated 30° relative to the animal’s dorsoventral axis and tilted 70° off the horizon to ensure that the stimulus was perpendicular to the optic axis of the eye (Oommen and Stahl, 2008). A behavior stage was placed in a consistent location using a kinematic mount and consisted of a standardized headframe clamp to enable repeatable positioning of the mouse relative to the monitor, and a 6.5” running wheel tilted upwards by 10-15 degrees. Running behavior was measured by a rotational encoder. Water rewards were delivered using a solenoid (NResearch, Model #161K011) that allowed for a calibrated volume of fluid to pass through a blunted, 17g hyperdermic needle (Hamilton) positioned approximately2-3mm from the animal’s mouth. Licks were detected by a capacitive sensor coupled to the reward delivery spout. Running speed, lick times, and reward delivery times were recorded on a NI PCI-6612 digital IO board and sampled at the frequency of the visual display (60 Hz).

#### Behavioral training procedure

Mice were trained for 1 hour/day, 5 days/week using an automated training algorithm. Briefly, mice were trained to lick when the identity of a flashed visual stimulus changed. If mice responded correctly within a short, post-change response window (750ms) a water reward (5-10uL) was delivered. On Day 1 of the automated training procedure mice received a short, 15-min “open loop” conditioning session during which non-contingent water rewards were delivered coincident with 90 degree changes in orientation of a full-field, static square-wave grating. This session was intended to 1) introduce the mouse to the fluid delivery system and, 2) provide the technician an opportunity to identify the optimal lick spout position for each mouse and 3) condition the association between stimulus changes and reward delivery. Each session thereafter was run in “closed loop” mode, and progressed through 3 stages of the operant task (schematized in Figure 1D): 1) static, full-field square wave gratings (changes between 0 and 90 degrees), 2) flashed, full-field square-wave gratings (changes between 0 and 90 degrees) presented for 250ms with an 500ms inter stimulus gray period, and 3) flashed full-field natural scenes (8 natural images from the Allen Brain Observatory) presented for 250ms with an 500ms inter stimulus gray period. Progression through each stage required mice to achieve a session maximum d’ of 2 on two of the last 3 sessions. Thus, the shortest amount of time to reach the final stage of training was 5 sessions. Once in stage 3, mice were considered ‘ready for imaging’ when 2 out of 3 sequential sessions had a d’ greater than 1 and mice performed at least 100 contingent trials. Mice were transitioned to behavior on the two-photon rig as scheduled time on the microscope became available. This resulted in a variable training duration in stage 3 across mice (Supplemental Figure 1A).

#### Session and trial structure

Each behavior session consisted of a continuous series of GO and CATCH trials, schematized in Figure 1B. Briefly, prior to the start of each trial a change-type and change-time were selected. Change-type was chosen based on predetermined frequencies such that GO and CATCH trials occurred with equal probabilities for sessions with 2 oriented gratings. For the natural image phase in which there were 64 change-pair possibilities, CATCH frequency was set to 12.5% (1/8 of the number of image transitions). To ensure even sampling of all stimulus transitions, a transition path is selected at random from a matrix of 1000 pre-generated paths. Each path takes a pre-determined route through each of the 64 possible transitions, including same-to-same, or catch, transitions. Once a transition path is completed, another path is chosen at random.

Change times were selected from an exponential distribution ranging from 2.25 to 8.25 seconds (mean of 4.25 seconds) following the start of a trial. Catch trial times were drawn from the same distribution such that false alarm rates were measured with the same temporal statistics as change trials, to account for any learning of the temporal distribution of change times. On trials when a mouse licked prior to the change or catch time, the trial was restarted with the same scheduled change or catch time. To prevent mice from getting stuck on a single trial, the number of times a trial could be repeated was limited to five. In all, this trial structure permits equal sampling of GO and CATCH trials, that when combined with mouse’s licking response, yields HIT, MISS, FALSE ALARM, and CORRECT REJECTION trials. In addition to the four trial types described above, behavior sessions contained a subset of “free reward” trials (GO trials followed immediately by delivery of a non-contingent reward). Behavior sessions across all phases began with 5 free-reward trials. Additionally, to promote continued task engagement, one of these free rewards was delivered after 10 consecutive MISS trials.

### Two photon imaging during behavior

#### Visual Stimulation

Visual stimuli were generated using custom Python scripts written in PsychoPy (Peirce, 2007, 2008) and were displayed using an ASUS PA248Q LCD monitor, with 1920 x 1200 pixels. Stimuli were presented monocularly, and the monitor was positioned 15 cm from the mouse’s eye, and spanned 120° X 95° of visual space. The monitor was rotated 30° relative to the animal’s midline and tilted 70° off the horizon to ensure that the stimulus was perpendicular to the optic axis of the eye (Oommen and Stahl, 2008).

The monitor was gamma corrected and had a mean luminance of 50 cd/m^2^. To account for the close viewing angle of the mouse, a spherical warping was applied to all stimuli to ensure that the apparent size, speed, and spatial frequency were constant across the monitor as seen from the mouse’s perspective (Marshel et al., 2011). Visual stimuli were presented at 60Hz frame rate.

Visual stimuli consisted of a subset of the natural scene images used in the publicly available Allen Brain Observatory dataset (https://observatory.brain-map.org/visualcoding/). The 32 natural images that we used originated from 3 different databases of natural scene images: the Berkeley Segmentation Dataset (images 000, 005, 012, 013, 024, 031, 034, 035, 036, 044, 047, 045, 054, 057) (Strasburger et al., 2011), the van Hateren Natural Image Dataset (images 061, 062, 063, 065, 066, 069, 072, 073, 075, 077, 078, 085, 087, 091) (van Hateren and van der Schaaf, 1998), and the McGill Calibrated Colour Image Database (images 104, 106, 114, 115) (Olmos and Kingdom, 2004). The images were presented in grayscale, contrast normalized, matched to have equal mean luminance, and resized to 1174 X 918 pixels.

#### Behavior apparatus

Running speed measurement, lick detection and reward delivery were performed as described above for behavioral training. The monitor was placed in a fixed location relative to the behavior stage to ensure a consistent relationship between the mouse’s eye and the screen. Running speed, lick times, and reward delivery times were recorded on a NI PCI-6612 digital IO board and sampled at the frequency of the visual display (60 Hz).

#### Two-photon calcium imaging during behavior

Calcium imaging was performed using a Scientifica Vivoscope (https://www.scientifica.uk.com/products/scientifica-vivoscope). Laser excitation was provided by a Ti:Sapphire laser (Chameleon Vision – Coherent) at 910 nm. Pre-compensation was set at −10,000 fs2. Movies were recorded at 30Hz using resonant scanners over a 400 μm field of view (512×512 pixels). Temporal synchronization of calcium imaging, visual stimulation, reward delivery and behavioral output (lick times and running speed) was achieved by recording all experimental clocks on a single NI PCI-6612 digital IO board at 100 kHz.

Behavior sessions under the two-photon microscope were 1 hour in duration, with task parameters identical to stage 3 of the behavior training procedure as described above. In addition, during most two-photon imaging sessions, 5% of stimulus flashes were randomly omitted, excluding the change flash and the flash immediately prior to the change. These omitted flashes were added to the experimental protocol partway into the experiment, resulting in 86/101 (85%) imaging sessions including omitted flashes. The 15 sessions without omitted flashes included data from one Slc17a7-IRES2;CaMKII-tTA;Ai93 mouse (4 sessions in VISp), and two Vip-IRES-Cre;Ai148 mice (3 sessions from VISal, and 8 sessions from VISp). Sessions without omitted flashes were excluded from any analysis depending on stimulus omission.

Movies of fluorescence were acquired near the center of retinotopic space in VISp and VISal, using ISI target maps and vasculature images as a guide. Once a cortical region was selected, the objective was shielded from stray light coming from the stimulus monitor using opaque black tape. All recordings were made at a depth of ∼175um from the brain surface. Once a field of view was selected, a combination of PMT gain and laser power was selected to maximize laser power (based on a look-up table against depth) and dynamic range while avoiding pixel saturation (max number of saturated pixels <1000). Immersion water was occasionally supplemented while imaging using a micropipette taped to the objective (Microfil MF28G67-5 WPI) and connected to a 5 ml syringe via an extension tubing. At the end of each experimental session, a z-stack of images (+/- 30 μm around imaging site, 0.1 μm step) was collected to evaluate cortical anatomy and evaluate z-drift during experiment. Experiments with z-drift above 10µm over the course of the entire session were excluded.

For each field of view, imaging and behavior sessions were conducted using each of the 4 image sets shown in Figure 1C, including the familiar image set A used during behavior training, and 3 novel image sets first experienced by the mouse during the imaging phase of the experiment. On subsequent imaging days for a given field of view, we returned to the same location by matching (1) the pattern of vessels in epi-fluorescence with (2) the pattern of vessels in two photon imaging and (3) the pattern of cellular labelling in two photon imaging at the previously recorded location. Typically, only one field of view was imaged per mouse, however in 3 out of the 21 mice, fields of view were recorded in both VISp and VISal. In cases where an imaging session failed our QC criteria (for example for z-drift >10um, or due to hardware issues such as dropped stimulus of imaging frames; see below), the session was retaken. As a result, some sessions with ‘novel’ image sets B, C or D were the second or third exposure (67% were first exposure, 27% were the second exposure, 6% were the third or fourth exposure). In contrast, mice were exposed to familiar image set A for an average of 17 +/-14 sessions during training.

### Quality control

All data streams were required to pass an initial integrity check. Frame sync times for 2-photon can have no more than 6 dropped frames, and a mean physiology period (frame rate) between 0.032 - 0.034. The visual stimulus presentation sync times can have no more than 60 dropped frames and an average frame interval between 0.0165 and 0.0167. The display lag of the monitor is measured using a photodiode to compare with recorded frame times, and the display lag must not exceed 150ms. The running wheel encoder data stream is examined for any visible artifacts (such as a spike in the trace or a flat trace despite running activity). The average intensity of the 2-photon field of view may not drift more than 10% over the course of a session. The acquired movie is checked for saturation to ensure that no more than 500 saturated pixels are present across the duration of the recording session. Z-drift is quantified by performing phase correlation between the frames of a 100um z-stack taken after the imaging session and a 500 frame average from the beginning of the 2-photon movie and a 500 frame average at the end of the movie. If the distance between the z-stack frames found to be most correlated with the beginning and end of the movie is greater than 10um, the session is retaken.

### Data processing

All data processing was performed as described in de Vries et al., 2018.

For each two-photon imaging session, the image processing pipeline included the following steps: 1) motion correction, 2) image normalization to minimize confounding random variations between sessions, 3) segmentation of ROIs, and 4) ROI filtering. Motion correction was performed using phase correlation and rigid translation. Segmentation was performed by morphological filtering on normalized periodic average images constructed from 400 frame blocks, followed by unification of masks across all blocks. ROI filtering was performed to remove segmented regions that were unlikely to correspond to cell somas, based on attributes including size and shape (for example, small ROIs likely corresponding to apical dendrites were removed).

Following identification of cell ROIs, the following steps were performed to obtain *ΔF/F* (dF/F) traces: 1) neuropil subtraction, 2) trace demixing, 3) *ΔF/F* computation. For each ROI, a neuropil mask was created, consisting of a 13 pixel ring around the cell soma, excluding any other ROIs. The raw fluorescence trace was generated by averaging all pixels within each cell ROI and the neuropil mask. A neuropil contamination ratio was computed for each ROI and the calcium trace was modeled as *F*_*M*_= *F*_*c*_+ *rF*_*N*_, where *F*_*M*_ is the measured fluorescence trace, *F*_*C*_ is the unknown true ROI fluorescence trace, *F*_*N*_ is the fluorescence of the surrounding neuropil, and *r* is the contamination ratio. After determination of *r*, we computed the true trace as *F*_*C*_ = *F*_*M*_− *rF*_*N*_, which is used in all subsequent analysis. To avoid artificially correlating neurons’ activity by averaging fluorescence over two spatially overlapping ROIs, we demixed the activity of all recorded ROIs, as described de Vries et al., 2018. Finally, the global dF/F trace for each cell was computed, with the baseline F_0_ determined by a rolling median filter of 180 seconds across the raw fluorescence trace.

Temporal alignment was performed to link two-photon acquisition frames (30Hz frame rate) with visual stimulation frames (60Hz frame rate) and associated behavioral signals (licking, running speed, reward delivery, sampled at 60Hz frame rate of visual stimulus). The visual stimulus time nearest to each two-photon (2P) frame time was computed, with the condition that the visual stimulus time must be before the 2P acquisition time, to ensure that dF/F responses were not attributed to stimulus or behavior events occurring after the change in the calcium signal.

### Data Analysis

#### Behavior

Response rates for GO and CATCH trials were calculated by evaluating the fraction of trials of each type where a lick was registered within the 750ms response window following the change or sham change time (Figure 1H). The fraction of GO trials with a response is the hit rate and the fraction of CATCH trials with a lick response is the false alarm rate. The d-prime value for each session was computed as:

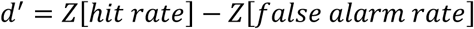

Where Z is the inverse of the cumulative distribution function (using scipy.stats.norm.ppf).

Reaction time was calculated as the time to first lick after the start of the change time on GO trials. Mean run speed was calculated by taking the average of the running speed trace in a +/-2 second window around the image change time for each GO trial, then averaging across all GO trials in each session (Supplemental Figure 1D).The average running speed trace across sessions (Supplemental Figure 1E) was computed by averaging the running speed trace across all GO trials in a [-2,6] second window around the change time for GO trials or the sham change time for CATCH trials.

Calculation of all behavior metrics was limited to the portion of the session where the mouse was actively engaged in the behavioral task, where engagement was defined those periods during which the mouse earned at least 2 rewards per minute. Mice performed 248 engaged GO trials per session on average (range = 83-335).

#### Physiology

All analysis was performed on the global dF/F traces (where baseline F was computed as the mode of a rolling 3-minute window, described in data processing section above).

Neural responses were analyzed for three main conditions of interest – the time of image presentations (across all stimulus flashes), the time of image changes (on GO trials, first flash after an image transition), and the time of stimulus omissions (5% of all non-change image flashes were randomly omitted). For each individual image presentation (or image omission), the mean response in a 500ms window after stimulus onset (or the time of omission) was computed (including the 250ms stimulus duration and 250ms after, to account for the slow decay of GCaMP6 responses and to include cells with delayed responses or off responses after stimulus offset).

The preferred stimulus for a cell was identified as the image evoking the largest trial-averaged response for a given condition. When considering all image presentations, the preferred stimulus is the image that evoked the largest mean response, averaged across all stimulus presentations of each image. For image changes, the preferred stimulus is the image that evoked the largest mean change response, considering only the first image presentation after a change in stimulus identity. Thus, an individual cell may have a different preferred image when considering all image flashes versus only the change flash. For some analyses of stimulus omissions, we considered omission trials where the preceding image was the cell’s preferred image defined across all stimulus presentations. Other analyses were agnostic to the identity of the image before and after the omission. Any analyses considering the image prior to the omission are indicated in text and figure legends.

Responsiveness was evaluated on a trial by trial basis (here an individual image presentation is considered as a ‘trial’). The mean response for each image flash was compared to a shuffled distribution of dF/F values taken from omission periods (the longest period of extended gray screen during the session) for each cell. A p-value was computed by resampling the shuffled distribution 10,000 times and determining the fraction of comparisons where the mean response was larger than the shuffled values. If the trial had a significantly larger response compared to the shuffled distribution, that trial was deemed responsive. For a cell to be considered *image responsive*, at least 25% of all stimulus presentations for the preferred image for that cell must be significant per the above definition (Figure 2G). The distribution of fraction significant image trials (for each cell’s preferred image) is shown in Supplemental Figure 2B. For a cell to be considered *change responsive*, at least 25% of all change flashes of the preferred image must be significant (Figure 2H). The distribution of fraction of significant change trials (for each cell’s preferred change image) is shown in Supplemental Figure 2C. For omission responsiveness, a different definition was used. In this case, a p-value was computed comparing the dF/F trace during the omission window (750ms after the omitted stimulus onset time, up to the time of the next stimulus onset) with a 750ms second baseline period prior to the omission (which includes the prior stimulus presentation and one inter-stimulus gray period). If 25% of all stimulus omission trials had a p-value > 0.05, with the mean omission response being greater than the baseline response, the cell was considered *omission responsive* (Figure 2I). The distribution of fraction of responsive omission trials is shown in Supplemental Figure 2C.

To evaluate image selectivity, we created tuning curves of mean response across the 8 images shown in each session, for each cell, for both the change flash (Supplemental Figure 3) and across all flashes (Figure 3). We generated a normalized image tuning curve for each image set by sorting each cell’s tuning curve by the strength of the mean response to each image, normalizing to the max, and averaging across cells (Figure 3G, Supplemental Figure 3E). Only cells meeting the criteria for image or change responsiveness described above were included in the calculation of the image tuning curve. An insufficient number (<10) of VIP cells were image responsive for image set A, thus a tuning curve was not included for this condition in Figure 3G. To quantify selectivity for individual cells, we used a lifetime sparseness metric, computed using the definition in (Vinje and Gallant, 2000):

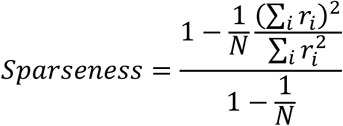

where *N* is the number of images and *r*_*i*_ is the response of the neuron to image i averaged across trials. Lifetime sparseness was only computed for image or change responsive cells. Population sparseness was computed with the same metric, but where N is the number of neurons and *r*_*i*_ is average response vector of neuron i to all images (Figure 3D, Supplemental Figure 3C). Lifetime and population sparseness were computed for the change flash only (Supplemental Figure 3), and for the mean response across all image flashes (Figure 3).

We computed a reliability metric by taking the average of the trial-to-trial cross correlation of the dF/F trace in a [-0.5, 0.75] second window around the stimulus onset time for the preferred image for each cell (see example cells in Figure 4D). This metric was evaluated under several conditions: for change responsive cells (Supplemental Figure 4E), for cells that were image selective (lifetime sparseness value > 0.3) vs. unselective (lifetime sparseness < 0.3) (Supplemental Figure 4F), and as a function of stimulus repetition number after a change for the preferred image (repetition 1 = first flash after a change, repetition = 10 is the 10^th^ flash in the sequence for that image) (Figure 4E).

To better understand the difference in response strength between the first flash after a change in image identity compared to after multiple repetitions of a given image, we computed a change modulation index (CMI):

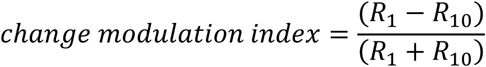

Where R_1_ is the trial averaged response to the 1^st^ flash after a change and R_10_ is the trial averaged response to the 10^th^ flash after a change. This metric was computed for each cell, either taking the trial average across all images (Supplemental Figure 4D) or the trial average only for each cell’s preferred image (Supplemental 4C) then averaged across all cells within a session to produce the plots in Supplemental Figures C&D. For Supplemental Figure 4B, the CMI listed in the title for each plot was computed using the mean of the population average trace across all images for the 1^st^ and 10^th^ repetitions.

A stimulus modulation index, measuring the modulation of the response at the stimulus frequency, was computed as in (Matteucci et al., 2019):

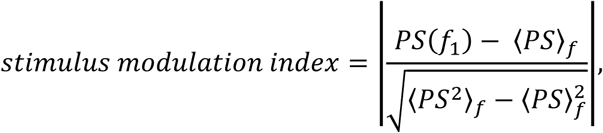

Where PS is the power spectral density (computed using scipy.signal.welch() with nperseg=128 and fs= 1/0.75) of the trial-averaged response over an 8 second window after the time of the change to the cell’s preferred stimulus (time = 0-8sec in example figures in Figure 4G), f_1_ is the stimulus frequency (1/0.75, images are presented every 750ms), and ⟨ ⟩_*f*_ indicates the average over frequencies. This metric quantifies the difference between the power at the stimulus frequency and the average of the power spectrum. The distribution of the signal modulation index value for the preferred image across cells is shown in Figure 4H.

The dynamics of cell responses were evaluated by computing a ramp index over different time windows of interest, similar to (Makino and Komiyama, 2015):

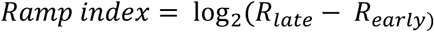

Where *R*_*late*)_ is the mean response in the first half of a defined window of time, and *R*_*early*)_ is the second half of the window. This index provides a measure of the magnitude and direction of a change in a signal within the window. For Figure 5, the ramp index was computed for two windows: the pre-stimulus window (250ms prior to stimulus onset; Figures 5A&B light shading) and the stimulus window (250ms after stimulus offset; Figures 5A&B dark shading) for the mean dF/F trace for each cell across all flashes of its preferred image. If the dF/F values are increasing during the window, the ramp index is positive. If the dF/F values are decreasing during the window, the ramp index is negative. The ramp index was only computed for cells with a mean dF/F value of >0.05 in the stimulus window, as the ramp index for a flat, unchanging signal can result in extreme values with little meaning.

The pre-stimulus and stimulus ramp indices were plotted against each other on a cell by cell basis (for cells with a minimum level of activity as described above) and found to be correlated by least squares linear regression between the two measures, performed separately for each image set (scipy.stats.linregress). Cells fell into four quadrants of this graph based on the relative sign of pre-stimulus and stimulus ramping. Cells with positive values of the stimulus ramp index and negative values of the pre-stimulus ramp index, indicating stimulus evoked activity with no pre-stimulus ramping, belonged to quadrant 1 (Q1). Most cells fell into this category. At the other extreme, cells with negative values of the stimulus ramp index and positive values of the pre-stimulus ramp index fall in Q4, indicating increasing activity prior to stimulus onset, and decreasing activity after stimulus offset. These cells showed the largest change in proportion between the familiar and novel image sets, particularly for VIP cells (Figure 5D). The fraction of cells belonging to each quadrant was computed for each imaging session, then averaged across sessions, in figure 5D. To illustrate the response dynamics associated with these cell groups, the mean response across all cells belonging to each quadrant was computed, then normalized to its max, for the plots in Figure 5E.

After grouping cells by their response dynamics, several metrics were computed, as described previously, for each group, separated by Cre line and image set, including: time to peak response for the average of all image flashes (Figure 5F), the stimulus modulation index (Figure 5G), reliability across all image flashes (Figure 5H), and lifetime sparseness across all flashes (Figure 5I). Only cells with a minimum level of activity (<0.05 dF/F) were included in this analysis. Plots in Figures 5F-I show the mean+/-95% confidence intervals across included cells.

The ramp index described above was again used to quantify the increase in activity during stimulus omission (Figure 6). For cells with a minimum level of activity during the omission period (>0.05 mean dF/F; Figure 6E), the ramp index was computed over the 750ms window, taking 250ms starting at the time where the omitted stimulus would have been presented as the early portion of the window, and the 250ms prior to the beginning of the next image presentation as the late portion of the window (note that 250ms in the middle of this 750ms window was unused). A positive value of this index indicates increasing activity over the omission period. The distribution of the ramp index values over this window across cells with a minimum level of activity during omission quantified in Figure 6F. The relationship between pre-stimulus ramping and omitted ramping is shown in Figure 5G on a cell by cell basis, with the strength and significance of the correlation determined using least squares linear regression (scipy.stats.linregress) as before. The fraction of cells having both increasing activity during the pre-stimulus period and increasing activity following stimulus omission are quantified in Figure 5H.

#### Statistics and data visualization

All statistical comparisons were made across image sets, within each cell class. ANOVA (scipy.stats.f_oneway) was used to test for an effect of image set, followed by a paired t-test (scipy.stats.ttest_ind) with Bonferroni correction for multiple comparisons for each individual image set pair. p-values are reported throughout the text and figure legends, and significance of comparisons (p<0.05) is indicated by an asterisks in figure insets. Point plots (seaborn.pointplot()) show individual sessions as gray points and the mean +/- 95% confidence intervals in color. Point plots lacking gray points are the mean +/- 95% confidence intervals across cells. Cumulative distributions across cells were generated with seaborn.distplot() with hist=False, hist_kws={‘cumulative’:True}. Barplot show either the mean across cells or the mean +/-SEM across cells. Heatmaps of cell responses were created using seaborn.heatmap().

